# Proteome changes associated with effect of high-dose single-fractionation radiation on lung adenocarcinoma cell lines

**DOI:** 10.1101/2025.01.21.634048

**Authors:** Elina Leis, Vijayachitra Modhukur, Helen Lust, Markus Vardja, Merilin Saarma, Ivar Ilves, Jana Jaal, Darja Lavogina

## Abstract

Lung cancer is a leading cause of cancer-related mortality globally, with non-small cell lung cancer (NSCLC) representing 85% of cases. Advances in treatment modalities, including the emergence of antibody-drug conjugates and stereotactic radiation therapy, have improved outcomes. However, the possible synergistic effects of these therapies remain underexplored at the molecular level. This study investigated high-dose radiation-induced proteomic changes in lung adenocarcinoma cell line HCC-44 grown adherently and cell line A549, grown as adherent cells and 3D spheroids. Our hypothesis was that proteins upregulated by 10 Gy irradiation serve as resistance drivers in cancerous cells and can thus represent potential therapeutic targets.

The label-free mass spectrometry revealed distinct proteomic responses to 10 Gy irradiation, varying by cell line and culturing conditions. Differentially expressed proteins elevated in the irradiated samples included ephrin type-A receptor 2 (EPHA2) in adherent cells and insulin-like growth factor 2 receptor (IGF2R), tetraspanin 3 (TSPAN3) as well as cathepsin D (CTSD) in spheroids. The validation of these targets was carried out *via* Western blot, immunofluorescence, viability assay and spheroid formation assay. The functional assays demonstrated that irradiation sensitized A549 cells to EPHA2 and CTSD inhibitors. These findings underscore the potential of integrating radiation and targeted therapies in NSCLC treatment, and highlight EPHA2 as a promising candidate for future therapeutic strategies.

## 1. Introduction

Lung cancer remains to be a leading cause of cancer-related deaths worldwide. There are two major types of lung cancers: small-cell lung cancer and non-small cell lung cancer (NSCLC), the latter comprising about 85% of all the lung cancer types. Historically, metastatic NSCLC (mNSCLC) has had a poor prognosis and 5-year overall survival (OS) has stayed under 10% (1). Over time, treatment options have evolved gradually, and combining different treatment modalities may improve survival significantly.

Oligometastatic disease (characterized by the presence of up to 5 metastases in up to 2 different organs) with its improved survival rates is now being distinguished from other metastatic diseases. In 2020, long-term results of the SABR-COMET Phase II randomized trial were made available. In this study, patients with oligometastatic disease were treated with either standard-of-care therapy (SOC) or SOC with radical local therapy using stereotactic radiation therapy; the 5-year OS rate was 17.7% *vs* 42.3%, respectively (2). Systemic therapy has also evolved over time and new pharmaceuticals have emerged to clinical practice, making treatment more effective while reducing the potential side effects. Antibody-drug conjugates (ADCs) represent a novel class anti-cancer drugs that bring much value to the treatment of various metastatic cancer sites, such as those in metastatic breast cancer and lung cancer (3,4). ADCs consist of three main components: antibody binding to the extracellular part of the target of interest, cytotoxic payload, and the connecting linker. In contrast to therapeutic monoclonal antibodies, the expression level of a specific target has less impact on the effectiveness of ADCs, where the cytotoxic payload plays a more significant role. For example, in the phase 3 ASCENT trial, metastatic triple negative breast cancer patients with low TROP2 expression still demonstrated improved progression-free survival with ADC sacituzumab govitecan (5).

Increasing number of ADCs in lung cancer treatment are undergoing clinical trials. Already established targets are human epidermal growth factor receptor 2 (HER2) with trastuzumab deruxtecan, HER3 with patritumab deruxtecan, trophoblast cell surface antigen 2 (TROP2) with sacituzumab govitecan, tyrosine kinase receptor c-Met (MET) with telisotuzumab vedotin. Currently in Europe, trastuzumab deruxtecan is the only approved ADC in second line NSCLC with HER2 activating mutations (6). There are also several early-phase clinical trials assessing additional targets-of-interest, for example NECTIN4, mesothelin, and ephrin receptor A2 (EphA2) (7,8). While combinations of systemic therapy and local therapy agents have been investigated intensely over time, there is still little clinically relevant information whether stereotactic radiation therapy enhances the effect of ADCs.

Here, we set out to explore the high-dose radiation-induced changes in the proteome of lung adenocarcinoma cell lines grown adherently or as 3D spheroids. The high-dose radiation approach was chosen as it imitates clinically relevant stereotactic radiation therapy. Using high-dose radiation in preclinical model mimics stereotactic radiation therapy in clinical setting. As the *in vitro* models, we chose two well-characterized cell lines that we have studied previously: HCC-44, expressing high levels of PD-L1 and featuring *KRAS* activating mutation G12C as well as *TP53* loss-of-function mutations (9); and A549, expressing low levels of PD-L1 and featuring *KRAS* activating mutation G12S and loss-of-function mutations in *STK11* and *KEAP1* (10). We hypothesized that the proteins upregulated by radiation will represent the survival strategies of cancerous cells, and the identified upregulated proteins could therefore serve as putative drug targets in future studies.

## 2. Materials and methods

### 2.1. Chemicals, cell lines and equipment

The list of materials used was mostly analogous to our previous studies (11–13). Human non-small cell lung carcinoma (adenocarcinoma) cell line HCC-44 and human lung carcinoma (adenocarcinoma) cell line A549 were from the Leibniz Institute DSMZ (German Collection of Microorganisms and Cell Cultures GmbH). The solutions and growth medium components for the cell culture were obtained from the following sources: phosphate-buffered saline (PBS), foetal bovine serum (FBS), L-glutamine, and Dulbecco’s Modified Eagle’s medium (DMEM) – Sigma-Aldrich (Steinheim, Germany); a mixture of penicillin, streptomycin, and amphotericin B – Capricorn (Ebsdorfergrund, Germany). For treatment of cells, ALW II-41-27 and pepstatin A (Selleckchem; Munich, Germany) were used.

The cells were grown at 37 °C in 5% CO2 humidified incubator (Sanyo; Osaka, Japan). The number of seeded or collected cells was counted using TC-10 cell counter (Bio-Rad; Hercules, CA, USA). During the sample treatment prior to proteomics or Western blot, the cells were grown either as adherent culture on the Falcon 75 cm² canted neck tissue culture-treated flasks with vented caps (Corning; Durham, North Carolina, USA) or as scaffold-free spheroids on Elplasia 12K flasks (Corning; Oneonta, New York, USA). For irradiation, the flasks were positioned between the 2 slabs of solid water phantom (30 cm × 30 cm × 5 cm below, 30 cm × 30 cm × 1 cm above the plates) and irradiated with dose 10 Gy (Gantry angle 0°, collimator angle 0°, field size 25 cm × 25 cm, source to surface distance 100 cm, dose rate 0.6 Gy/min). During the radiation studies, the cells were exposed to 6 MV X-rays (Varian Truebeam 2.5). For microscopy studies, the cells were seeded onto 24-well tissue culture-treated Ibidi black μ-plates (ibidi GmbH, Gräfelfing, Germany). In case of the viability assay, the cells were seeded onto transparent 96-well clear flat bottom cell culture plates BioLite 130188 (Thermo Fischer Scientific, Rochester, NY, USA). For transfer of the cells to irradiation facility, gas-permeable adhesive moisture seals were used (Brooks Life Sciences, Wotton, Surrey, UK). In case of spheroid formation assay, 96-well black ultra-low attachment spheroid microplates with clear round bottom were utilized (Corning 4515; Kennebunk, ME, USA).

For proteomics, dithiothreitol (DTT) was purchased from VWR Life Science (Radnor, PA, USA), chloroacetamide (CAA), ammonium bicarbonate (ABC), methylamine, guanidine hydrochloride (Gu-HCl), urea and thiourea from Sigma Aldrich (St. Louis, MI, USA). All chemicals were of proteomics grade or ≥99% purity. Lys-C and dimethylated trypsin used for protein digestion were purchased from New England Biolabs (Ipswich, MA, USA) and Sigma Aldrich (St. Louis, MI, USA), respectively. All organic solvents used for proteomics were of LC/MS grade from Honeywell (Charlotte, NC, USA).

The liquid chromatography with tandem mass spectrometry (LC/MS/MS) apparatus consisted of a Dionex (Sunnyvale, CA, USA) Ultimate 3000 RSLCnano chromatography system coupled to a Thermo Fisher Scientific (Waltham, MA, USA) Q Exactive HF mass spectrometer. The nano-LC setup consisted of a Dionex cartridge pre-column (ID 0.3 mm × L 5 mm, 5 μm C18) and a MS Wil (Aarle-Rixtel, the Netherlands) emitter-column (ID 75 μm × L 50 cm) packed with 3 μm C18 particles (Dr Maisch, Ammerbuch, Germany).

For lysis of cells prior to Western blot, HEPES and NaCl from Calbiochem (Darmstadt, Germany), Triton X from Ferak (Berlin, Germany), EDTA-containing cOmplete™ protease inhibitor cocktail from Roche (Basel, Switzerland) and PMSF from AppliChem (Darmstadt, Germany) were used. The total protein content of lysates was established using Pierce™ Coomassie Plus (Bradford) Assay Reagent (Thermo Fischer Scientific; Rockford, IL, USA) and the absorbance was measured at 590 nm using PHERAstar multi-mode reader (BMG Labtech; Ortenberg, Germany). For preparation of SDS-PAGE samples, 4× NuPAGE™ LDS Sample Buffer (Thermo Fischer Scientific; Carlsbad, CA, USA) supplemented with 50 mM DTT (Darmstadt, Germany) was used. Protein transfer was performed onto the PVDF membrane (Roche; Mannheim, Germany) in NuPAGE™ transfer buffer (Thermo Fischer Scientific; Carlsbad, CA, USA) supplemented with methanol (Honeywell, Riedel-de Haën™, Seelze, Germany). For the preparation of blocking solution used in Western blot and immunofluorescence (IF) experiments, BSA was obtained from Capricorn Scientific (Ebsdorfergrund, Germany) and PBS (supplemented with Ca^2+^, Mg^2+^) from Sigma-Aldrich (Steinheim, Germany). The rabbit monoclonal antibody against human Ephrin Type-A Receptor 2 (anti-EPHA2, clone D4A2) was from Cell Signaling Technology (catalogue number #6997; Danvers, Massachusetts, USA); the rabbit monoclonal antibody against human Insulin-Like Growth Factor 2 Receptor (anti-IGF2R) was from Sigma-Aldrich (catalogue number HPA011332; Saint Louis, Missouri, USA); the mouse monoclonal antibody against Tubulin Alpha 1a (anti-TUBA1A, clone DM1A) was from Novus Biologicals (catalogue number NB100-690; Centennial, Colorado, USA). As the secondary antibodies, alkaline phosphatase-conjugated secondary antibodies from Thermo Fischer Scientific (goat anti-rabbit T2191 or goat anti-mouse T2192; Bedford, MA, USA) were used in case of Western blot.

In case of IF, the same primary antibodies were used as in case of Western blot; additionally, the rabbit monoclonal antibody against human Tetraspanin 3 (anti-TSPAN3) was purchased from Sigma-Aldrich (catalogue number HPA015996; Saint Louis, Missouri, USA). The secondary antibodies used in IF [goat cross-adsorbed antibody against rabbit IgG (H+L), conjugated with Alexa Fluor 568; goat cross-adsorbed antibody against mouse IgG (H+L), conjugated with Alexa Fluor 647] and the nuclear stain 4’,6-diamidino-2-phenylindole (DAPI) were from Invitrogen (Eugene, Oregon, USA). Fluorescence microscopy with immunostained cells was carried out with Cytation 5 multi-mode reader using 20× air objective (0.3225 µm/pixel). For DAPI, 365 nm LED and DAPI filter block were used; for Alexa Fluor 568, 523 nm LED and RFP filter block were used; for Alexa Fluor 647, 628 nm LED and CY5 filter block were used.

Resazurin and PBS for viability assay (supplemented with Ca^2+^, Mg^2+^) were from Sigma-Aldrich (St Louis, MO, USA). Imaging of spheroids was carried out with Cytation 5 multi-mode reader using bright-field microscopy with 4× air objective (1.613 µm/pixel) and automated focussing regime.

### 2.2. Cell treatment prior to mass-spectrometry

In case of adherent culturing, HCC-44 or A549 cells (passage number below 15) were seeded in growth medium (DMEM supplemented with 10% FBS) onto the 75 cm² flasks (at 1/6 dilution from a confluent flask) and grown for 48 h as in culture. Next, irradiation of some of the flasks was carried out at 10 Gy. After 72 h post-irradiation, the spent culture media were collected into centrifuge tubes; the cells were rinsed with PBS, detached from the plates using 0.25% trypsin, resuspended in the culture medium, and then combined with the corresponding spent media aliquots to collect both detached dying cells and the surviving population. 30 μL aliquots of the obtained cell suspension (total volume of 9 mL per flask) were taken for counting of non-disintegrated cells (the results are shown in Supplementary Figure S1).

In case of 3D culturing, A549 cells (passage number below 15) were seeded in growth medium onto the Elplasia 12K flasks (at 1/5 dilution from a confluent flask) and grown for 48 h as in culture. Next, irradiation of some of the flasks was carried out at 10 Gy. After 72 h post-irradiation, the spent culture media were collected into centrifuge tubes. The flasks were rinsed with the growth media and PBS and all solutions were combined with the corresponding spent media aliquots. HCC-44 cells were not used in these experiments as we could not establish a protocol for obtaining compact spheroids from this cell line.

The following treatment was performed according to the procedures established previously (13). The cells or spheroids were pelleted from the collected suspensions by centrifugation (5 min at 800 rcf) and the pellets were washed twice with PBS. Finally, PBS was removed, and dry pellets were frozen and stored at −90 °C until all independent experiments (N = 3) were finished.

After transportation on dry ice to the proteomics facility, the pellets were suspended in 10 volumes of 6 M Gu-HCl, 100 mM Tris-HCl pH 8.5, 50 mM DTT lysis buffer. Samples were heated at 95 °C for 5 min, followed by Bioruptor sonication (Diagenode) for 15 min (30 s on, 60 s off, 10 cycles) at “high” setting at 4 °C. *Ca* 1 volume of 0.5 mm Zirconia/Silica beads (BioSpec Products) was added to the lysate, which was then homogenised in a FastPrep24 (MP Biomedicals) device using 2× 40 s 6 m/s pulses. Sample tubes were punctured with a heated needle and the solution was centrifuged from the beads to a new 2.0 ml LoBind tube at 2000 rpm for 2 min. Non-soluble material was pelleted by centrifugation at 17,000 g for 10 min at 4 °C. For the full proteome analysis, 10 µg of protein was precipitated with trichloroacetic acid/deoxycholate. Protein pellets were suspended in 20 µL of 7 M urea, 2 M thiourea, 100 mM ABC, 2 mM methylamine solution, followed by disulphide reduction and cysteine alkylation with 10 mM DTT for 30 min at 30 °C and 30 mM CAA for 30 min at room temperature in the dark. Proteins were pre-digested with 1:100 (enzyme to protein) Lys-C for 2 h, diluted 5 times with 100 mM ABC and further digested with 1:50 trypsin overnight at 25 °C. Peptides were desalted with in-house made C18 StageTips (14) and reconstituted in 0.5% trifluoroacetic acid.

### 2.3. Label-free proteomics

The methodology used was analogous to our previous study (13). 2 µg of peptides were injected onto a 0.3 × 5 mm trap-column (5 µm C18 particles, Dionex), from which they were eluted to an in-house packed (3 µm C18 particles, Dr Maisch) analytical 50 cm × 75 µm emitter-column MS Wil (Aarle-Rixtel, the Netherlands). The columns were operated at 45 °C. Peptides were separated at 300 nL/min with an 8-40% A to B 120 min gradient. Eluent B was 80% acetonitrile + 0.1% formic acid and eluent A was 0.1% formic acid in water. Eluted peptides were injected into a Q Exactive HF hybrid quadrupole – Orbitrap (Thermo Fisher Scientific) tandem mass spectrometer using a nano-electrospray source and a spray voltage of 2.5 kV (liquid junction connection). The MS instrument was operated with a top-12 data-dependent acquisition strategy. One 350-1400 m/z MS scan (at a resolution setting of 60 000 at 200 m/z) was followed by a MS/MS (R=30 000 at 200 m/z) of the 12 most intense ions using higher-energy collisional dissociation fragmentation (normalized collision energy of 26). The MS and MS/MS ion target and injection time values were 3×10^6^ (50 ms) and 1×10^5^ (41 ms), respectively. The dynamic exclusion time was limited to 50 s; only charge states +2 to +5 were subjected to MS/MS.

### 2.4. Bioinformatic analysis of proteomic data

The analysis pipeline was analogous to our previous study (13). MS raw data were processed with the MaxQuant software (version 2.2.0.0) (15), searching for the protein and peptide matches from the UniProt human reference proteome database (downloaded 09.21.2023) (16). Methionine oxidation, and protein N-terminal acetylation were set as variable modifications, while cysteine carbamidomethylation was defined as a fixed modification. Tryptic digestion rule (cleavages after lysine and arginine without proline restriction) was used for *in silico* digestion of the database. Only identifications with at least 1 peptide ≥ 7 amino acids long (with up to 2 missed cleavages) were accepted and transfer of identifications between runs based on accurate mass and retention time (‘match between runs’) was enabled. Label-free normalization with MaxQuant LFQ algorithm was also applied. Protein and LFQ ratio count (*i.e*., number of quantified peptides for reporting a protein intensity) was set to 1. Peptide-spectrum match and protein false discovery rate was kept below 1% using a target-decoy approach. All other parameters were default.

Statistical software R v4.2.3 and package DEP (17) were used for downstream analysis of the quantified proteins. During preprocessing, proteins that were identified in less than two out of three replicates in at least one condition were filtered out. Next, the proteome data was background corrected and normalized by variance stabilizing transformation. To account for missing values, the k-nearest neighbour imputation method was applied. Principal component analysis (PCA) was performed and plotted using the plot_pca function from the DEP package to visualize sample variance and clustering between conditions. Prior to differential analysis in irradiated *versus* non-irradiated cells, counts were logarithm-transformed. R package limma was used for differential analysis (irradiated *versus* non-irradiated cells in different cell lines and culturing conditions) based on the linear models and empirical Bayes statistics (18). Proteins were considered significantly different at FDR < 0.05. The final DEP lists (Supplementary Table S1) were generated by eliminating proteins for which the razor + unique peptide count in two out of three replicates considered for the differential analysis was less than 2. Volcano plots were generated using the plot_volcano function from the DEP package to visualize the protein fold changes (x-axis) and adjusted p-values (y-axis) between the compared conditions. The final DEP lists were analysed using the g:Profiler online platform (version e111_eg58_p18_f463989d, database updated on 25/01/2024 (19)) to identify the significantly altered pathways associated with the radiation treatment (Supplementary Table S2).

### 2.5. Western blot

The treatment of the cells (2D-cultured HCC-44, 2D-cultured A549, A549 spheroids) and collection and storage of the cell pellets was carried out in the same way as described for the mass-spectrometry studies. The lysis of cells was carried out as previously described (20). For preparation of SDS-PAGE samples, the total protein concentrations in lysate supernatants were established using Bradford assay and the concentrations of samples were adjusted according to the most dilute sample. The SDS-PAGE samples were denatured under reducing conditions by 15 min incubation at 75 °C; prior to SDS-PAGE, the samples were stored at −20 °C. The subsequent procedures were carried out according to the previously published protocol (20) with exception of the transfer time (25 min at 20 V was used in this study).

### 2.6. Immunostaining and fluorescence microscopy

The cells were seeded onto the 24-well plates with the working volume of 0.5 mL and density of 4000 or 8000 cells per well (for HCC-44 and A549, respectively) and grown overnight. Next, irradiation of cells at 10 Gy was carried out as described above; the non-irradiated cells were kept on a separate plate. To prevent achieving overly high density of the cells on the non-irradiated plate, the cells were subsequently placed to the incubator and grown on the plate only for 48 h post-irradiation. The medium was removed, the cells were rinsed with PBS and fixed directly on the plate with cold methanol (15 min at −20 °C). Afterwards, methanol was removed, the cells were washed twice with PBS, and blocking with 1% BSA in PBS (weight/volume) was performed for 1 h at rt, followed by the staining and wash procedures analogously to the previously reported protocol (21) with the solution volumes adjusted for a 24-well plate. The following dilutions were used for the primary antibodies: in case of anti-EPHA2, 1:500; in case of anti-IGF2R, 1:400; in case of anti-TSPAN3, 1:200; and in case of anti-TUBA1A, 1:2000. The secondary antibodies were used at 1:1000 dilution; for the staining of nuclei, 300 nM DAPI in PBS was applied. The imaging parameters (LED intensity, signal integration time, and camera gain) were first optimized in the manual imaging mode for each antibody, and the same parameters were then used for this antibody for all plates in all independent experiments (N = 4). The imaging was performed in the automated mode; 25 images per well were taken and the DAPI channel was used for autofocusing.

### 2.7. Viability assay

The approach used was analogous to our previous studies (12,22). HCC-44 or A549 cells (passage number below 15) were seeded in growth medium onto the 96-well plate with the density of 2000 or 3500 cells per well, respectively (within the linear range of the method, optimized in previous studies (11)). The cells were left to attach for 24 h at 37 °C under the usual culturing conditions. Next, irradiation of some plates at 10 Gy was carried out. 2 h post-irradiation, the growth medium was exchanged, and 5-fold dilution series of biologically active compounds in growth medium were added onto the cells. Based on the solubility of compounds in the water, the maximal final total concentrations of compounds were 10 μM. An identical volume of growth medium was added to the negative control (100% viability). The final volume per well was 150 μL, and the concentration of DMSO in the treated wells was ≤ 0.1% by volume; on each plate, each concentration of each compound was represented in triplicate. To prevent achieving overly high density of the cells on the non-irradiated plate, the cells were subsequently placed to the incubator and grown on the plate only for 48 h post-irradiation. Next, and viability assay was then carried out according to the previously published protocol (11).

### 2.8. Spheroid formation assay

The approach used was analogous to our previous study (12). A549 cells (passage number below 15) were grown on the two 75 cm² flasks to 80% confluency. Next, irradiation of one flask at 10 Gy was carried out. 2 h post-irradiation, the cells were detached from the plates using 0.25% trypsin, resuspended in the culture medium and then seeded in growth medium onto the 96-well ultra-low attachment plate with the density of 4000 (experiment 1) or 2000 (experiments 2-3) cells per well. At the same time, 5-fold dilution series of compounds in the growth medium were added to the wells. The final total volume in the well was 200 μL and the maximal final total concentrations of compounds were 20 μM (as in general, higher concentrations of compounds are required in 3D cell culture as compared to 2D to achieve the same extent of biological effect). The plates were placed into the incubator and imaged at 72 h and 96 h post-seeding.

### 2.9. Other statistical analyses

The statistical analyses were carried out analogously to the previously reported procedures (11–13). Throughout the study, the grouped comparisons were carried out using non-parametric Kruskal-Wallis test with Dunn’s test for multiple comparisons; unless indicated otherwise, the pairwise comparisons were carried out using the unpaired two-tailed t-test with Welch’s correction. In all statistical tests, the significance of comparisons is indicated as follows: *** indicates P ≤ 0.001, ** indicates P ≤ 0.01, * indicates P ≤ 0.05.

In case of Western blot, the band intensities corresponding to the EPHA2 (110-130 kDa) or IGF2R (270 kDa) were normalized to the band intensity of the loading control (TUBA1A, 50 kDa) in each independent experiment (N = 4). Subsequently, additional normalization to the non-irradiated samples was then carried out separately for adherent HCC-44, adherent A549, or A549 spheroids (normalized ratio set to 100%). The normalized data for the identically treated cells was then pooled.

In case of IF, three frames without any visible artefacts (*e.g*., dust particles) were randomly chosen within a set of 25 frames per well. The mean signal intensity in RFP channel (corresponding to the antibody of interest) was quantified in three representative regions of interest per frame (each region of interest involving 1-3 individual cells) using raw images with non-modified brightness or contrast. The data for the identically treated cells was then pooled for all independent experiments (N = 4). Non-parametric Mann-Whitney test was used for the paired comparisons as the analysed data did not follow the normal distribution according to the D’Agostino and Pearson omnibus normality test.

In case of viability assay, in each independent experiment, the fluorescence intensity measured for the replicate treatments was pooled and the data obtained for the negative control was plotted against incubation time with resazurin. One time-point within duration of data acquisition was chosen where the signal of the negative control remained in the linear range, and only data measured at this time-point was used for the further analysis. For normalization, data obtained for wells treated with PBS (blank control) was considered as 100% viability; data acquired for the 50 μM resazurin solution (in the absence of cells) was considered as 0% viability. Next, the ratio of absorbance at 570 nm and 600 nm was calculated for each well. The ratios were analysed analogously to the fluorescence intensity data, and the normalized viability values calculated from the fluorescence intensity and the absorbance measurements were pooled. Finally, data from all independent experiments (N = 3 for each treatment in each cell line with 3 technical replicates per experiment) was pooled for each individual compound concentration. The pooled normalized viability was plotted against the concentration of compound in the dilution series and fitted to the biphasic equation with the Hill slope values fixed at −1 and the fraction of the curve derived from the more potent phase fixed between 0 and 1.

In spheroid formation assay, the spheroid contour was denoted manually using the freehand selections tool and the area and circularity of the spheroid were quantified; in case of disintegrated spheroids with poorly defined borders, the selection involved all dark area covered with cells. The spheroid area was quantified separately for the different time-points in each independent experiment (N = 3 for each treatment with 2 technical replicates per experiment) and normalized by the negative control (cell not treated with inhibitor or radiation) in each independent experiment (area set to 100%). The normalized data for the identically treated cells was then pooled.

### 2.10. Other software

For general data analysis, GraphPad Prism 6 (San Diego, CA, USA) and Excel 2016 (Microsoft Office 365; Redmond, WA, USA) were used. To create the Venn diagram, we utilized the interactive tool Venny (23). In case of Western blot, IF and spheroid formation assay, ImageJ software (Fiji package (24)) was used for the image analysis.

## 3. Results

### 3.1. Proteome changes associated with irradiation are dependent on the cell line and the mode of culturing

As the first step of our study, we performed mass-spectrometric profiling of the proteome corresponding to the non-irradiated or 10 Gy-irradiated cells grown adherently (in case of both HCC-44 and A549) or as spheroids (only in case of A549). The cells were left to recover for 72 h after irradiation and three independent experiments were carried out for each treatment.

We identified a total of 5188 proteins (present in at least one of the measured samples); in each individual sample, the number of detected proteins exceeded 4000, with minor variation between the treatments and independent experiments (see Supplementary Figure S2). This indicated that the chosen timing was correct, avoiding excessive cell death in the irradiated samples and thus enabling characterization a sufficiently versatile proteome. According to the PCA plot (Figure 1A), the clusters corresponding to the A549 vs HCC-44 cell line were separated along the PC1 dimension and the clusters corresponding to the adherent vs spheroid A549 cells were separated along the PC2 dimension. The difference between the clusters corresponding to the irradiated vs non-irradiated cells was less pronounced, yet still distinguishable for both cell lines and culturing conditions in case of A549 cells.

**Figure 1.**
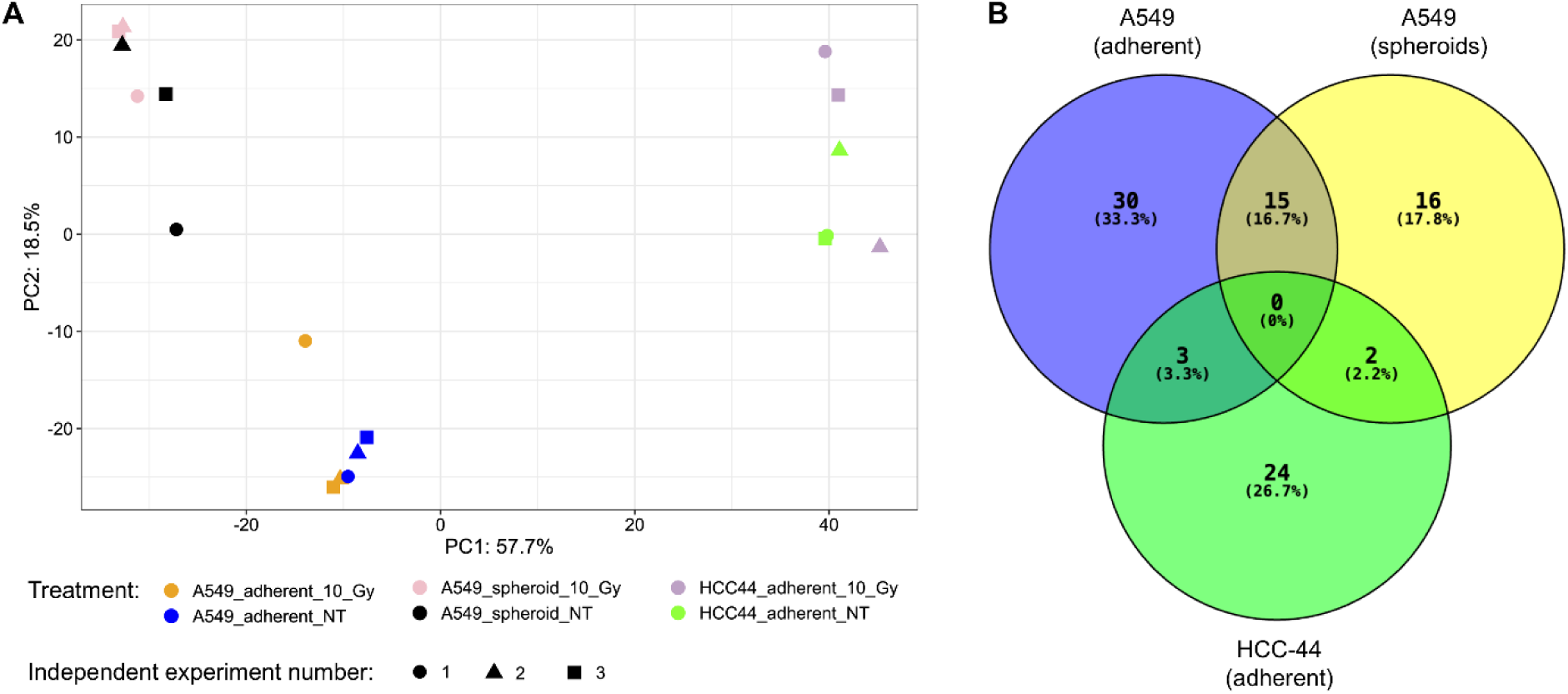
Characteristic clustering and differentially expressed proteins according to the mass-spectrometry data. (A) PCA plot of the proteomic data indicating differentiation between the treatments (N = 3 for each treatment). The plot was generated using the complete mass-spectrometry data after imputation. The colour codes for different cell lines, culturing modes and irradiation conditions are shown below the plot; NT corresponds to not irradiated cells. (B) Venn diagram of the DEPs in irradiated *vs* non-irradiated cells in different cell lines and culturing conditions (FDR < 0.05 in any compared treatments).

Next, we established the proteins which were significantly up- or downregulated by the irradiation (FDR < 0.05). The Volcano plots corresponding to three compared categories (adherent HCC-44 cells, adherent A549 cells and A549 spheroids) are presented in Supplementary Figures S3-S5. The number of differentially expressed proteins (DEPs, see Supplementary Table S1) was highest in the adherent A549 cells (48 DEPs, of which 26 were upregulated and 22 downregulated by irradiation), followed by the A549 spheroids (33 DEPs, of which 12 were upregulated and 21 downregulated by irradiation) and the adherent HCC-44 cells (29 DEPs, of which 22 were upregulated and 7 downregulated by irradiation). According to the Venn diagram (Figure 1B), the overlap of DEPs was highest in case of A549 adherent cells and A549 spheroids (15 DEPs). Only three DEPs were shared between the adherent A549 cells and HCC-44 cells [ephrin type-A receptor 2 (EPHA2), optineurin (OPTN) and microtubule-associated protein 1B (MAP1B)], and only two DEPs between A549 spheroids and adherent HCC-44 cells [aldehyde dehydrogenase 1 family member A3 (ALDH1A3) and karyopherin subunit alpha 2 (KPNA2)]. No common DEPs were found between all three compared categories.

The further analysis of DEPs focussed separately on the upregulated and downregulated proteins in each compared category. Based on the DEPs, we identified the significantly upregulated or downregulated pathways using the online platform g:Profiler; the results are presented in Supplementary Table S2 and summarized concisely in Table 1. Overall, the pathway analysis confirmed that HCC-44 cell line responded to irradiation differently than A549 cell line, while A549 adherent cells and spheroids shared multiple downregulated but not upregulated pathways. Particularly, in HCC-44, upregulation of G2/M markers and downregulation of multiple metabolic processes occurred upon irradiation. In the adherent A549 cells, the upregulated pathways were rather related to cytoskeleton and cellular adhesion, while downregulated pathways involved DEPs responsible for the DNA damage response. The latter observation was also valid in the context of A549 spheroids, while no significantly upregulated pathways could be identified in this model. The generally known functions related to the DEPs upregulated in irradiated spheroids involved the acyl-coenzyme A signalling [represented by the acyl-CoA dehydrogenase family member 8 (ACAD8) and acyl-CoA thioesterase 8 (ACOT8)], oxidoreductive catalysis [represented by ACAD8, ferredoxin reductase (FDXR) and 5-oxoprolinase (OPLAH)], and vesicle formation and trafficking [represented by cathepsin D (CTSD), DnaJ homolog subfamily C member 13 (DNAJC13), insulin-like growth factor 2 receptor (IGF2R), OPLAH and tetraspanin 3 (TSPAN3)].

**Table 1.** Examples of the differentially expressed proteins and the corresponding significantly enriched pathways, cellular locations and processes identified by g:Profiler in irradiated *versus* non-irradiated cells.

| Cell line and culture type | Comparison | Examples of DEPs | Examples of pathways, locations and processes |
| --- | --- | --- | --- |
| **HCC-44, adherent** | Upregulated in irradiated cells | KIF23, OPTN, PLA2G4A, TUBB2A, TUBB4A, TUBB6 | • intra-Golgi and retrograde Golgi-to-ER traffic<br>• mitotic spindle<br>• G2/M transition |
|  | Downregulated in irradiated cells | ABCC3, CAT, EPHX1, SCP2, SELENBP1 | glucuronoside transmembrane transporter activity<br>methanethiol oxidase activity<br>propanoyl-CoA C-acyltransferase activity<br>peroxisomal matrix<br>photodynamic therapy induced NFE2L2 NRF2 survival signaling |
| **A549, adherent** | Upregulated in irradiated cells | EPHA2, KRT18, MAP1B, MYH9, PALLD, PLEC, RAI14, TGFBI, TRIOBP | actin binding<br>cell adhesion molecule binding<br>stress fiber |
|  | Downregulated in irradiated cells | ADNP, MCM2, MCM3, MCM4, MCM5, MCM7, MSH2, MSH6, NASP, NCAPG, NUMA1, POLD3, RBBP7, RRM1, SMC2, SMC3, TOP2A, WDHD1 | catalytic activity, acting on DNA<br>DNA binding<br>cell cycle<br>DNA damage response<br>chromosomal region |
| **A549, spheroids** | Upregulated in irradiated cells | ACAD8, ACOT8, CTSD, DNAJC13, FDXR, IGF2R, OPLAH, TSPAN3 | None identified |
|  | Downregulated in irradiated cells | DNMT1, MCM2, MCM3, MCM4, MCM5, MCM6, MCM7, MSH6, NCAPD2, PCNA, PDS5A, POLD1, RRM1, SMC2, SMC4, TMPO, TOP2A, TRIP13 | catalytic activity, acting on DNA<br>DNA binding<br>cell cycle<br>DNA damage response<br>chromosomal region |

Abbreviations: DEP, differentially expressed protein

Based on the mass-spectrometry results, we chose for the further validation three proteins for which the membranous localization has been reported previously (25–27): EPHA2, which was elevated upon irradiation in both HCC-44 and A549 cell lines grown adherently [binary logarithm of fold change (LFC) values of 1.1 and 0.72, respectively]; and IGF2R and TSPAN3, which were elevated upon irradiation in A549 spheroids (LFC values of 1.1 and 0.89, respectively).

### 3.2. Immunostaining confirms radiation-induced increase in EPHA2, IFG2R and TSPAN3 levels

Initially, we set out to confirm whether the radiation-induced increase in EPHA2 and IGF2R levels can be detected by Western blot in lysates of HCC-44 and A549 cells grown adherently, or in lysates of A549 spheroids. The treatment and the subsequent culturing of cells was carried out as in case of the proteomic experiment, with lysis performed at 72 h after irradiation.

As the membranous fraction of lysates had too high viscosity for carrying out the SDS-PAGE, only lysate supernatants were utilized for the Western blot. Representative examples of EPHA2 and IGF2R staining and pooled data summarizing the loading control- and radiation status-normalized immunostaining intensity are presented in Figure 2A-C; the images of full membranes are shown in Supplementary Figures S6-S7. According to Western blot, EPHA2 and IGF2R could be detected in both irradiated and non-irradiated cells. While the EPHA2 signal showed the trend towards increase upon irradiation in all compared categories (adherent HCC-44 cells, adherent A549 cells, and A549 spheroids), the increase was not statistically significant due to high variation between the experiments; in case of HCC-44, borderline significance was achieved (P < 0.1). In case of IGF2R, a trend towards increase upon irradiation was evident for adherent HCC-44 cells and A549 spheroids, whereas the latter trend was also statistically significant (P < 0.05).

**Figure 2.**
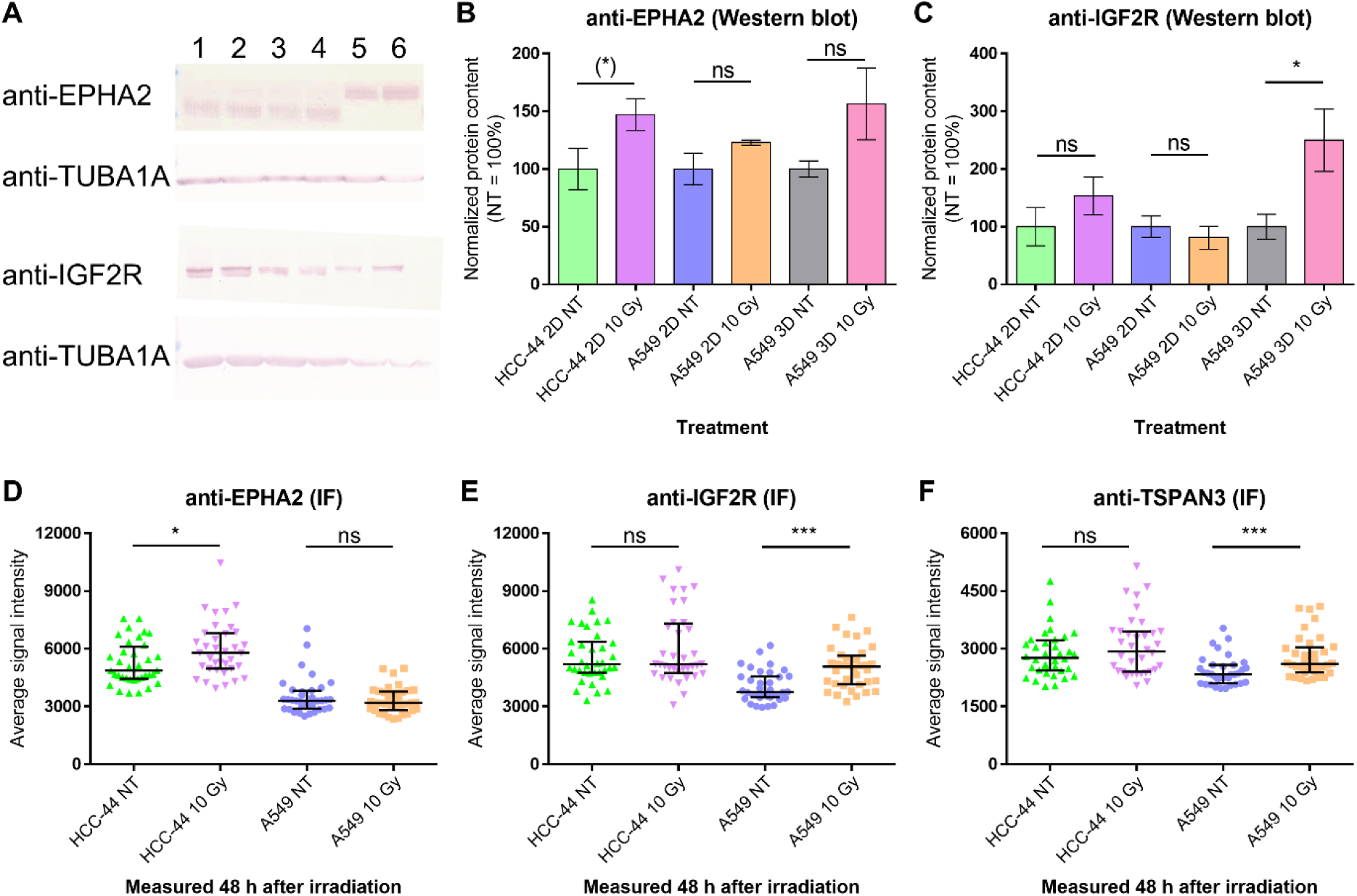
Validation of the radiation-induced expression of the targets chosen based on the proteomics data using Western blot (A-C) and immunofluorescence (D-F). (A) representatives examples of staining in Western blot; data from a single experiment with anti-EPHA2 and a single experiment with anti-IGF2R is shown. TUBA1A was used as the loading control; lane 1 corresponds to non-irradiated HCC-44 cells grown adherently, lane 2 to irradiated HCC-44 cells grown adherently, lane 3 to non-irradiated A549 cells grown adherently, lane 4 to irradiated A549 cells grown adherently, lane 5 to non-irradiated A549 cells grown as spheroids, lane 6 to irradiated A549 cells grown as spheroids. For better visualization, the contrast of image parts showing EPHA2 and IGF2R staining was reduced; the quantification shown in B and C was carried out using non-modified images. (B, C) Pooled Western blot staining data normalized first to loading control and then separately to the corresponding non-irradiated control (NT) in each independent experiment (N = 4). Each column represents average ± standard error of mean; the immunostained protein names are mentioned in the graph header and the cell lines and treatment conditions below the graph. Statistical significance of paired comparisons (t-test): * corresponds to P < 0.05; (*) corresponds to P < 0.1; ns, not significant. (D-E) Signal intensity established from the microscopy images following IF; the immunostained protein names are mentioned in the graph header and cell treatment conditions prior to lysis below the graph. Each point corresponds to one region of interest in one frame; thick black line shows median and whiskers show interquartile range; data from all independent experiments was pooled (N = 4). Statistical significance of paired comparisons (Mann-Whitney test): *** corresponds to P < 0.001; * corresponds to P < 0.05; ns, not significant.

Next, we explored the intensity of immunostaining of EPHA2, IFG2R and TSPAN3 in the adherently cultured HCC-44 and A549 cells by comparing the non-irradiated cells and cells fixed at 48 h after 10 Gy irradiation. The representative examples of microscopy images are shown in Figure 3 (EPHA2 staining) and Supplementary Figures S8-S9 (IGF2R, TSPAN3 staining); the pooled quantified signal intensity (N = 4) is presented in Figure 2D-F. Based on the microscopy images, all three proteins were detectable in both irradiated and non-irradiated samples. For EPHA2, the membranous and cytoplasmic staining was evident in both HCC-44 and A549 cells, with clear enrichment in mitotic cell population in both cell lines. On the other hand, IGF2R and TSPAN3 rather featured cytoplasmic and nuclear staining, while the membranous pool could not be confirmed for these proteins due to the overall weak staining intensity. During quantification of the staining intensity, we did not distinguish between the various intracellular localization. According to the pooled IF data, increase of EPHA2 expression upon irradiation was statistically significant in HCC-44 (P < 0.05) but not in A549 cells, while IGF2R and TSPAN3 were elevated upon irradiation only in A549 cells (P < 0.001). Overall, the trends observed in both Western blot and IF generally confirmed the mass-spectrometry results, although some cell line-dependent variation was evident.

**Figure 3.**
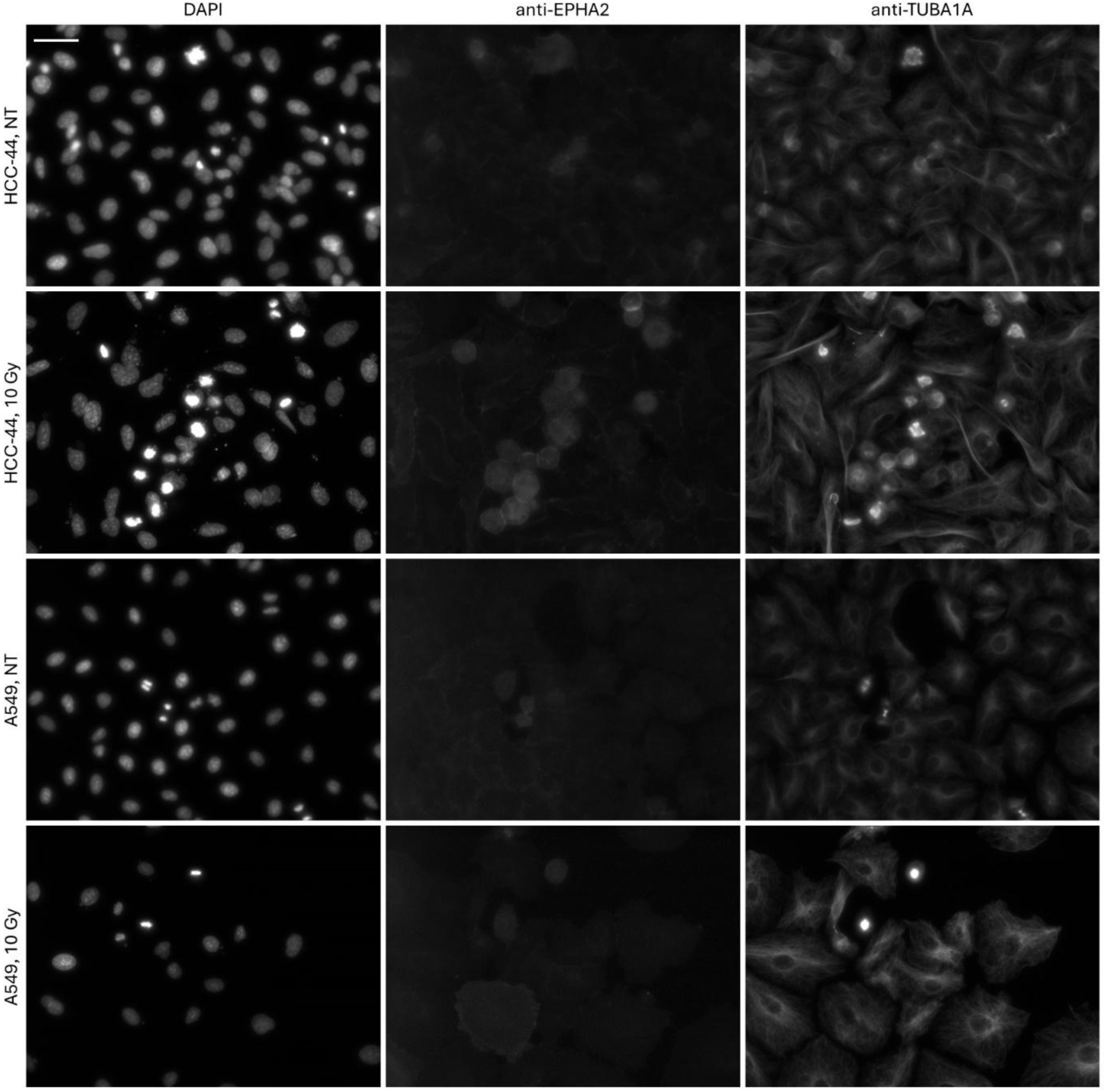
Representative examples of EPHA2 staining in irradiated *vs* non-irradiated adherent fixed cells. Data from a single experiment is shown; the channels (nuclear stain DAPI, protein of interest, and TUBA1A) are listed above the images; the cell lines and treatment conditions are listed on the left (NT stands for not irradiated). Scale bar (top left): 50 μm.

### 3.3. Radiation sensitizes A549 cell line to EPHA2 inhibition

Next, we decided to explore whether interfering with activity of the selected target proteins can affect the growth and proliferation rate of irradiated or non-irradiated lung adenocarcinoma cells. From the three validated proteins mentioned above, a selective inhibitor (ALW-II-41-27) was commercially available for EPHA2 (28). Furthermore, we expanded our validation set of targets with CTSD, which belongs under the aspartic protease family and for which an enzyme family-selective inhibitor pepstatin A was commercially available (29). According to our mass-spectrometry data, CTSD was elevated upon irradiation in A549 cell line independently on the mode of culturing (LFC values of 0.82 and 0.67 in adherent cells and spheroids, respectively; Supplementary Table S1).

We first assessed the viability/proliferation of the adherent HCC-44 and A549 cells using the resazurin assay at 48 h post-irradiation at 10 Gy; the inhibitors were added onto the cells shortly after irradiation. The results of resazurin assay are summarized in Figure 4; the normalization of data was carried out separately for the irradiated and non-irradiated cells in each independent experiment to enable better comparison between the putative changes in the dose-response profile. The dose-response profile followed the biphasic pattern for both compounds, which is generally indicative of different action mode of compounds at low *vs* high dose of inhibitor (11). In case of ALW-II-41-27, the low-dose IC50 value was in the subnanomolar range in both cell lines, while the high-dose IC50 value in the submicromolar range in HCC-44 cells and in a single-digit micromolar range in A549 cells. In case of pepstatin A, the low-dose IC50 value was also in the subnanomolar range in both cell lines, while the high-dose IC50 value was in the submillimolar range. Importantly, irradiation of A549 cells triggered sensitization to both inhibitors, which was evident from the significantly increased proportion of the low-dose effect in the corresponding dose-response curves (P < 0.01 for ALW-II-41-27 and P < 0.001 for pepstatin A); no such effect was evident for HCC-44 cells.

**Figure 4.**
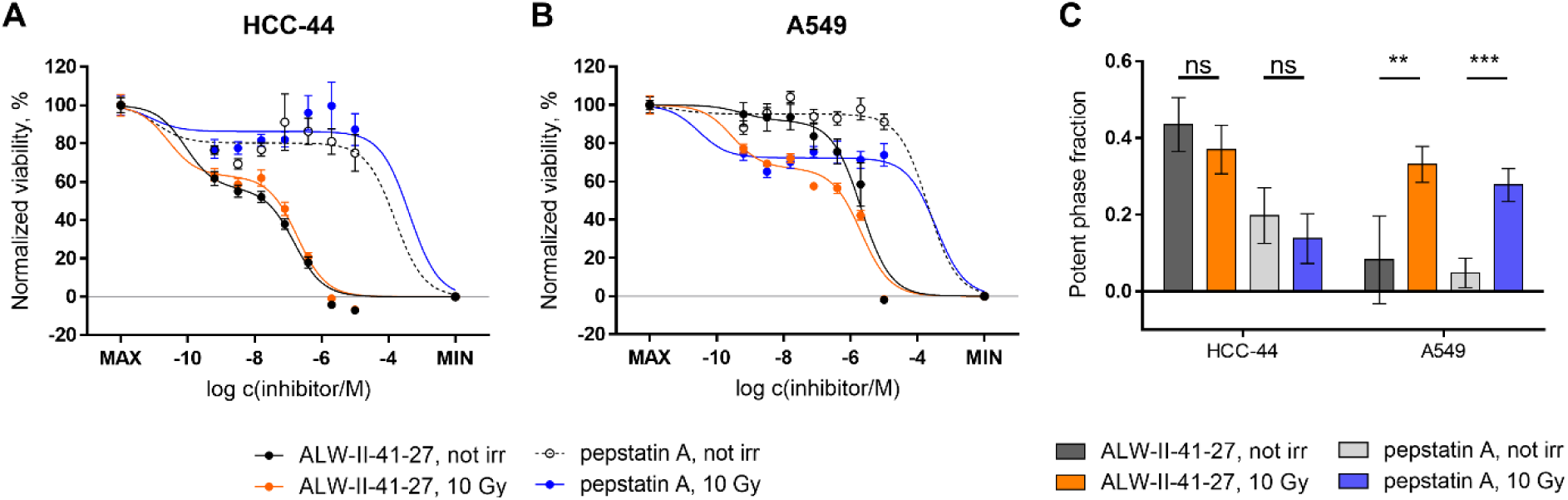
Viability/proliferation assay in irradiated *vs* non-irradiated adherent cells in the presence *vs* absence of EPHA2 and CTSD inhibitors. (A, B) Dose-response curves. Pooled normalized viability/proliferation is shown (N = 3); the cell line is indicated above the graphs and the treatment conditions below the graphs. Each column represents average ± standard error of mean; the data points were fit to the biphasic equation. MAX corresponds to the non-treated cells in case of non-irradiated cell dose-response and to 10 Gy-treated cells in case of the irradiated cell dose response. (C) Summary of the potent phase fraction statistics obtained based on the curves shown in panels A and B. The cell lines are indicated in the bottom part of the graph; each column represents average ± standard deviation. Statistical significance of the paired comparisons (t-test): *** corresponds to P < 0.001; ** corresponds to P < 0.01; ns, not significant.

We next explored whether the same inhibitors could interfere with the formation of spheroids in A549 cell line. For that, we seeded onto the ULA plate either 10 Gy-preirradiated or non-irradiated cells shortly after irradiation and added simultaneously the dilution series of either ALW-II-41-27 or pepstatin A onto the cells. After 72 h and 96 h post-seeding, the area of the formed spheroids was assessed by bright-field microscopy; a set of representative images is shown in Figure 5A-H and the pooled normalized results are summarized in Supplementary Figure S10 (72 h post-seeding) and Figure 5I (96 h post-seeding). In case of spheroid formation assay, the normalization of spheroid area was carried out according to the non-treated control only in each independent experiment to confirm the effect of irradiation on the spheroid size.

**Figure 5.**
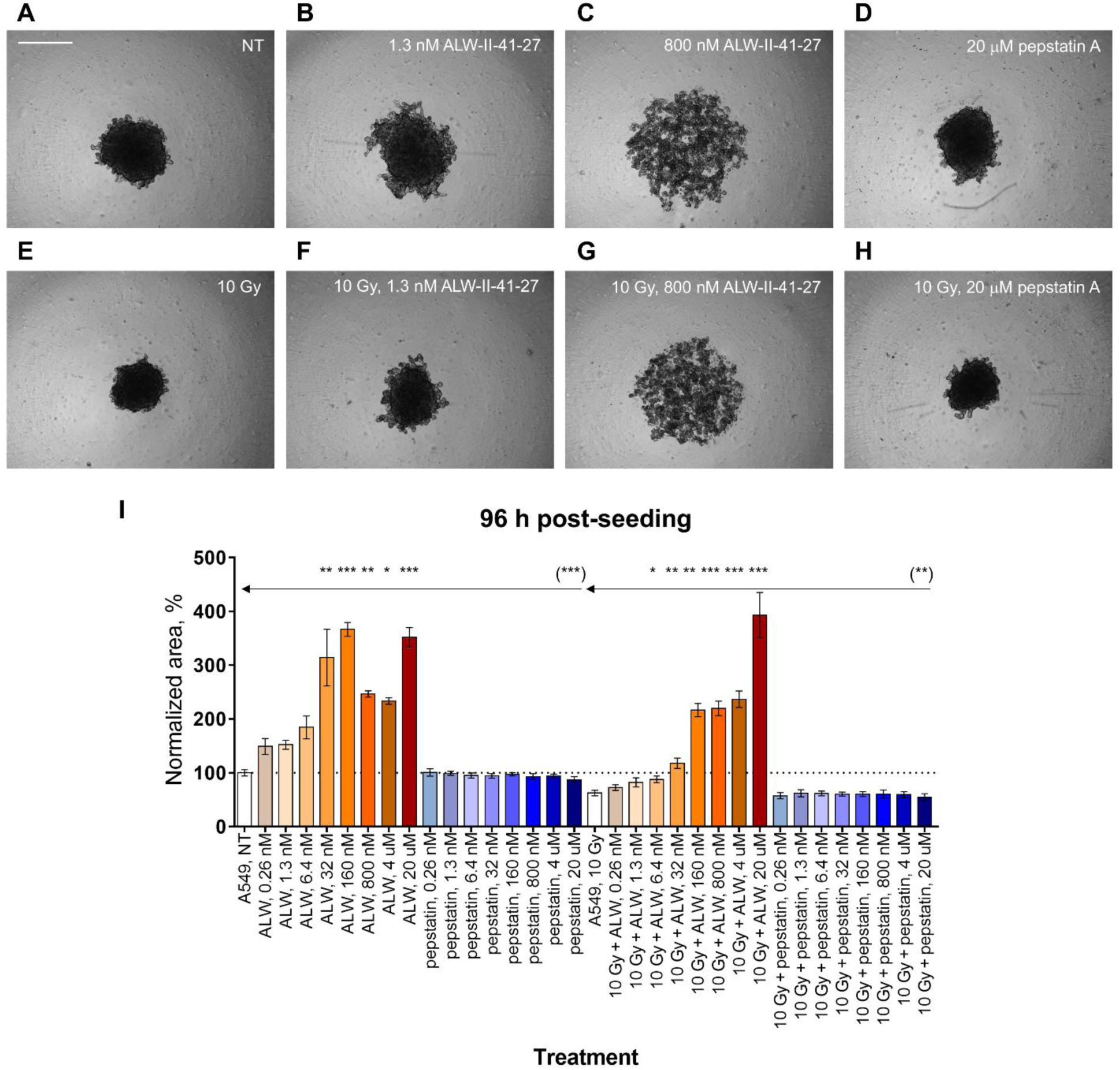
Spheroid formation assay in irradiated *vs* non-irradiated A549 cells in the presence *vs* absence of EPHA2 and CTSD inhibitors. (A-H) Representative examples of spheroids formed during 96 h post-seeding. Data from a single experiment is shown; the treatment conditions are listed in the top right corner of each panel (NT stands for not treated). Scale bar (top left): 400 μm. (I) Pooled normalized area of spheroids at 96 h post-seeding (N = 3); the treatment conditions are listed below the graphs. Each column represents average ± standard deviation. Statistical significance of the grouped comparisons (Kruskal-Wallis test): *** corresponds to P < 0.001; ** corresponds to P < 0.01; * corresponds to P < 0.05; only significant comparisons are shown; arrows indicate the treatments (non-treated A549 or irradiated A549) to which other treatments are compared. Statistical significance of paired comparisons (Mann-Whitney test, only carried out for pepstatin A treatment *versus* cells not treated with inhibitors): (***) corresponds to P < 0.001; (**) corresponds to P < 0.01; only significant comparisons are shown. Abbreviations: ALW, ALW-II-41-27; pepstatin, pepstatin A.

The spheroid formation assay confirmed the potency of the EPHA2 inhibitor ALW-II-41-27, which interfered with cell-cell contacts causing disassembly of spheroids at nanomolar concentrations of the compound. In irradiated cells, significant increase in spheroid size was evident at 6.4 nM concentration of ALW-II-41-27 (P < 0.05, multiple comparisons to the non-treated A549 cells), while in the non-irradiated cells, statistical significance became evident at 32 nM concentration of the compound (P < 0.01, multiple comparisons to the 10 Gy-irradiated A549 cells). On the other hand, pepstatin A caused significant reduction in spheroid size only at the highest concentration of the inhibitor tested (20 μM) irrespective of the irradiation status of the cells (P < 0.001 for the paired comparison of the non-irradiated A549 cells and P < 0.01 for the paired comparison of the irradiated A549 cells).

## 4. Discussion

Metastatic NSCLC treatment has significantly improved over time being shaped by the emergence of novel pharmaceuticals, such as targeted therapy and immunotherapy. During recent years, ADCs have shown impressive therapeutic activity against metastatic NSCLC and expanding targets as well as indications. Combining stereotactic radiation therapy with systemic therapy in the oligometastatic and oligoprogressive settings has also improved overall survival significantly (2). Whether combining novel-class pharmaceuticals ADCs with high-dose irradiation would also improve the outcome is yet to be investigated. This work is focused on finding post-irradiation proteomic changes in the hopes of finding new targets for ADCs and investigates whether the combination of ADCs and high dose irradiation has potential in preclinical models.

For our study, we chose lung adenocarcinoma cell lines A549 and HCC-44, which have been characterized in detail in previous studies by us (11,13,22) and numerous other research groups. Importantly, we have previously demonstrated that A549 cells are less sensitive to radiation as compared to HCC-44 cells: the former featured no reduction in viability at 48 h post-irradiation with 10 Gy and only 23% reduction in viability at 72 h post-irradiation with 8 Gy, while the latter featured correspondingly 21% and 48% reduction at the same conditions (11). Therefore, it was of special interest to investigate which molecular targets ensured post-irradiation survival of A549 cell line.

The proteomic changes at 72 h post-irradiation with 10 Gy were explored using the label-free proteomics strategy in lysates of both cell lines, analogously to our previous study with chemotherapeutic drug cisplatin where we had shown the presence of markers characteristic to either surviving or dying populations of cells (13). In the current study, the proteomes corresponding to different treatments clustered separately on the PCA plot (Figure 1A), being thus dependent on the cell line, the culturing method, as well as the irradiation status. The short-listing of DEPs (Supplementary Table S1) showed that in adherent cells, a higher number of upregulated proteins could be identified after irradiation as compared to the spheroids, which showed in turn a higher number of downregulated proteins upon irradiation (although this observation was limited by the fact that the spheroids could only be obtained for A549 cell line).

Based on the pathway analysis of the DEP lists (Table 1), it was evident that the adherent HCC-44 cells responded to irradiation by the cell cycle arrest in G2/M phase (*e.g*., upregulation of kinesin-like protein KIF23 (30) or karyopherin KPNA2 (31)). This was indicative of the DNA damage, thus being in line with the higher radiation sensitivity (32). Still, a population of HCC-44 cells also featured increase of adhesion marker EPHA2, which has been shown to contribute to cancer resistance mechanisms in NSCLC after irradiation (33). In turn, the adherent A549 responded to irradiation by the decreased damage response markers (*e.g*., cell-cycle-related minichromosome maintenance proteins MCM2-MCM6 (34) or condensing complex component SMC4 (35)) yet increased markers of cell adhesion (*e.g*., EPHA2 (36), non-muscle myosin MYH9 (37) or myoneurin PALLD (38)). While the decrease in MCM2-MCM6 cannot be unequivocally interpreted as the sign of resistance to therapy (as both overexpression and decrease in MCM protein levels has been shown to contribute to cancer (39)), the absence of clear cell cycle arrest markers and the increase of the cell-cell contact-ensuring proteins point to the cell adhesion-mediated radioresistance (40). Interestingly, several DEPs associated with radiation effect in this study were identical to the markers identified in our previous study where the same cell lines were grown adherently and treated with 1 μM cisplatin for 48 h (13): *e.g*., upregulation of KIF23, KPNA2 and EPHA2 in HCC-44, upregulation of ferredoxin reductase FDXR in A549, and downregulation of MCM2 in A549 was observed in both studies.

By contrast, A549 spheroids did not show a certain pattern of upregulated pathways in the current study (Table 1), likely mirroring the presence of several cell populations in spheroids consistent with the more physiological-like status but also complexity of the 3D-*versus* 2D-cultures. The pathways downregulated by irradiation were, however, similar in the adherently cultured A549 cells and spheroids, thus confirming the intrinsic tendency towards increased radioresistance of this cell line.

For validation, we chose EphA2, IGF2R, TSPAN3, and CTSD as these were elevated upon irradiation in at least one cell line or culturing condition according to the proteomic data (Supplementary Table S1). The three former proteins have been shown to localize on the cell plasma membrane (25–27) and could thus serve as putative targets for the antibody-drug conjugates in combination with irradiation as a future therapeutic strategy. Additionally, we decided to explore CTSD, which has been characterized as both lysosomal and extracellular protein (41), and for which inhibitor pepstatin A was commercially available. Furthermore, according to the literature, overexpression of IGF2R and CTSD can lead to radioresitance (42,43). Less evidence is available for TSPAN3, although it belongs to a family of tetraspanins than have been shown to regulate epithelial-mesenchymal transition, exosome signalling and other pathways sustaining survival of cells in various cancers including NSCLC (44,45). In our work, slight yet statistically significant elevation of both IGF2R and TSPAN3 levels upon irradiation was evident in adherent A549 by IF (P < 0.001 for both targets), and minor increase of IGF2R in the irradiated A549 spheroids was observed by Western blot (P < 0.05; Figure 2). These findings were in line with proteomics where IGF2R and TSPAN3 were found significantly elevated in irradiated A549 spheroids (FDR < 0.05) and showed a trend towards higher levels in irradiated *vs* non-irradiated adherent A549 (LFC value of 0.241 for IGF2R and LFC value of 0.908 for TSPAN3), although the statistical significance was not achieved for the adherent A549.

The good agreement of proteomics results with functional assays was further confirmed during validation of EphA2 as the radiation-induced target. Ephrin receptors constitute the largest family of the receptor tyrosine kinases and are directly affected by the cell-cell contacts as the activating ligands (ephrins) are themselves tethered to the membranes of the cells (46). In normal conditions, EphA2 is almost exclusively expressed in proliferating epithelial cells (47), yet in the pathological context, it is abundant in different cancer sites, including lung cancer. While EphA2 overexpression has been correlated to poor prognosis (48) and resistance to irradiation (33,49), there are several studies that report overcoming radioresistance by inhibiting EphA2 kinase activity or reducing EphA2 expression (28,50,51). Therefore, EphA2 is explored as a target in various novel anti-cancer therapies, including antibody-drug conjugates (8).

According to our assays, reliance of irradiated NSCLC cells on the EphA2-mediated pathways was confirmed in the context of all tested systems – adherent HCC-44 (higher levels of EphA2 in irradiated cells in Western blot and in IF; Figure 2), adherent A549 (higher proportion of the EphA2 inhibitor low-dose effect in irradiated cells; Figure 4), and A549 spheroids (significant effect of EphA2 inhibitor on spheroid area already at single-digit nanomolar concentration in pre-irradiated cells; Figure 5). While proteomics data indicated significant increase in EphA2 levels in adherent cells (FDR < 0.05), a trend towards higher levels in irradiated *vs* non-irradiated A549 spheroids was also observed (LFC value of 0.356), although the statistical significance was not achieved. The low nanomolar activity of EphA2 inhibitor ALW-II-41-27 was somewhat unexpected as it has been used at a remarkably higher (1 μM) concentration in studies of cervical cancer or NSCLC cell lines (28,52). The previous reports, however, did not explore either high-dose radiation of 3D cell culture context, which was addressed in detail in the current study.

This study has several limitations, including the use of only two lung adenocarcinoma cell lines, with one yielding self-assembling spheroids, and the validation of a limited number of targets using a single antibody or inhibitor per target. Additionally, variations in timing after irradiation were required for different validation methods to avoid excessive confluency in negative control cells. Despite these constraints, our findings confirm EPHA2 as a promising target for combating the surviving cancer cell population following irradiation. Also, because we chose more aggressive type on NSCLC cell line A549 with its low PD-L1 expression level, KRAS-activating mutation and loss-of-function mutations in STK11 and KEAP1, our results bring valuable information and new potential treatment options for future studies. Our study also especially with treatment resistant NSCLC. Overall, this study emphasizes the need for further clinical exploration of EPHA2. Moreover, other proteins of interest highlighted in this study should be assessed in patient biopsy specimens to provide valuable insights into their clinical relevance and role in cancer recurrence.

## Acknowledgements

We thank Ago Rinken group members for the maintenance of the microplate reader and microscopy platform.

## Data availability

The mass spectrometry proteomics data have been deposited to the ProteomeXchange Consortium via the PRIDE partner repository (53) with the dataset identifier PXD057576. Other datasets generated and analysed during the current study are available from the corresponding authors on reasonable request.

## Author roles

Conceptualization, E.L., J.J. and D.L.; Methodology, M.V., I.I. and D.L.; Validation, E.L., H.L., M.S. and D.L; Formal Analysis, V.M. and D.L.; Resources, I.I., J.J. and D.L.; Writing – Original Draft Preparation, E.L., V.M. and D.L.; Writing – Review & Editing, all authors; Visualization, V.M. and D.L.; Supervision, J.J and D.L.; Project Administration, J.J.; Funding Acquisition, J.J.

## References

1. Simeone JC, Nordstrom BL, Patel K, Klein AB. Treatment patterns and overall survival in metastatic non-small-cell lung cancer in a real-world, US setting. Future Oncol. 2019 Oct;15(30):3491–502.

2. Palma DA, Olson R, Harrow S, Gaede S, Louie AV, Haasbeek C, et al. Stereotactic Ablative Radiotherapy for the Comprehensive Treatment of Oligometastatic Cancers: Long-Term Results of the SABR-COMET Phase II Randomized Trial. J Clin Oncol. 2020 Sep 1;38(25):2830–8.

3. Bian DJH, Cohen SF, Lazaratos AM, Bouganim N, Dankner M. Antibody–Drug Conjugates for the Treatment of Non-Small Cell Lung Cancer with Central Nervous System Metastases. Current Oncology. 2024 Oct;31(10):6314–42.

4. Koster KL, Huober J, Joerger M. New antibody-drug conjugates (ADCs) in breast cancer-an overview of ADCs recently approved and in later stages of development. Explor Target Antitumor Ther. 2022;3(1):27–36.

5. Bardia A, Hurvitz SA, Tolaney SM, Loirat D, Punie K, Oliveira M, et al. Sacituzumab Govitecan in Metastatic Triple-Negative Breast Cancer. New England Journal of Medicine. 2021 Apr 21;384(16):1529–41.

6. Wass RE, Lang D, Horner A, Lamprecht B. Antibody–drug conjugates (ADCs) in lung cancer treatment. memo. 2024 Sep 1;17(3):198–203.

7. Coleman N, Yap TA, Heymach JV, Meric-Bernstam F, Le X. Antibody-drug conjugates in lung cancer: dawn of a new era? npj Precis Onc. 2023 Jan 11;7(1):1–12.

8. Xiao T, Xiao Y, Wang W, Tang YY, Xiao Z, Su M. Targeting EphA2 in cancer. Journal of Hematology & Oncology. 2020 Aug 18;13(1):114.

9. Ma’ayan Laboratory of Computational Systems Biology. COSMIC Cell Line Gene Mutation Profiles. [cited 2024 Oct 11]. Gene Set - HCC-44. Available from: https://maayanlab.cloud/Harmonizome/gene_set/HCC-44/COSMIC+Cell+Line+Gene+Mutation+Profiles

10. Ma’ayan Laboratory of Computational Systems Biology. COSMIC Cell Line Gene Mutation Profiles. [cited 2024 Oct 11]. Gene Set - A549. Available from: https://maayanlab.cloud/Harmonizome/gene_set/A549/COSMIC+Cell+Line+Gene+Mutation+Profiles

11. Lavogina D, Lust H, Tahk MJ, Laasfeld T, Vellama H, Nasirova N, et al. Revisiting the Resazurin-Based Sensing of Cellular Viability: Widening the Application Horizon. Biosensors (Basel). 2022 Mar 25;12(4):196.

12. Lavogina D, Krõlov MK, Vellama H, Modhukur V, Di Nisio V, Lust H, et al. Inhibition of epigenetic and cell cycle-related targets in glioblastoma cell lines reveals that onametostat reduces proliferation and viability in both normoxic and hypoxic conditions. Sci Rep. 2024 Feb 21;14(1):4303.

13. Saar M, Jaal J, Meltsov A, Laasfeld T, Lust H, Kasvandik S, et al. Exploring the Molecular Players behind the Potentiation of Chemotherapy Effects by Durvalumab in Lung Adenocarcinoma Cell Lines. Pharmaceutics. 2023 May 12;15(5):1485.

14. Rappsilber J, Mann M, Ishihama Y. Protocol for micro-purification, enrichment, pre-fractionation and storage of peptides for proteomics using StageTips. Nat Protoc. 2007 Aug;2(8):1896–906.

15. Tyanova S, Temu T, Cox J. The MaxQuant computational platform for mass spectrometry-based shotgun proteomics. Nat Protoc. 2016 Dec;11(12):2301–19.

16. The UniProt Consortium. UniProt: the Universal Protein Knowledgebase in 2023. Nucleic Acids Research. 2023 Jan 6;51(D1):D523–31.

17. Proteome-wide identification of ubiquitin interactions using UbIA-MS | Nature Protocols [Internet]. [cited 2024 Aug 19]. Available from: https://www.nature.com/articles/nprot.2017.147

18. Ritchie ME, Phipson B, Wu D, Hu Y, Law CW, Shi W, et al. limma powers differential expression analyses for RNA-sequencing and microarray studies. Nucleic Acids Res. 2015 Apr 20;43(7):e47.

19. Kolberg L, Raudvere U, Kuzmin I, Adler P, Vilo J, Peterson H. g:Profiler—interoperable web service for functional enrichment analysis and gene identifier mapping (2023 update). Nucleic Acids Research. 2023 Jul 5;51(W1):W207–12.

20. Sõrmus T, Lavogina D, Teearu A, Enkvist E, Uri A, Viht K. Construction of Covalent Bisubstrate Inhibitor of Protein Kinase Reacting with Cysteine Residue at Substrate-Binding Site. J Med Chem. 2022 Aug 25;65(16):10975–91.

21. Lavogina D, Laasfeld T, Vardja M, Lust H, Jaal J. Viability fingerprint of glioblastoma cell lines: roles of mitotic, proliferative, and epigenetic targets. Sci Rep. 2021 Oct 13;11(1):20338.

22. Saar M, Lavogina D, Lust H, Tamm H, Jaal J. Immune checkpoint inhibitors modulate the cytotoxic effect of chemotherapy in lung adenocarcinoma cells. Oncol Lett. 2023 Apr;25(4):152.

23. Oliveiros, J. C. Venny. An interactive tool for comparing lists with Venn’s diagrams. 2007 [cited 2024 Aug 19]. Venny 2.1.0. Available from: https://bioinfogp.cnb.csic.es/tools/venny/index.html

24. Schindelin J, Arganda-Carreras I, Frise E, Kaynig V, Longair M, Pietzsch T, et al. Fiji: an open-source platform for biological-image analysis. Nat Methods. 2012 Jul;9(7):676–82.

25. Chavent M, Seiradake E, Jones EY, Sansom MSP. Structures of the EphA2 Receptor at the Membrane: Role of Lipid Interactions. Structure. 2016 Feb 2;24(2):337–47.

26. Rezgui D, Williams C, Savage SA, Prince SN, Zaccheo OJ, Jones EY, et al. Structure and function of the human Gly1619Arg polymorphism of M6P/IGF2R domain 11 implicated in IGF2 dependent growth. J Mol Endocrinol. 2009 Apr;42(4):341–56.

27. Thiede-Stan NK, Tews B, Albrecht D, Ristic Z, Ewers H, Schwab ME. Tetraspanin-3 is an organizer of the multi-subunit Nogo-A signaling complex. J Cell Sci. 2015 Oct 1;128(19):3583–96.

28. Li X, Li D, Ma R. ALW-II-41-27, an EphA2 inhibitor, inhibits proliferation, migration and invasion of cervical cancer cells via inhibition of the RhoA/ROCK pathway. Oncol Lett. 2022 Apr;23(4):129.

29. Kozak A, Mikhaylov G, Khodakivskyi P, Goun E, Turk B, Vasiljeva O. A New Cathepsin D Targeting Drug Delivery System Based on Immunoliposomes Functionalized with Lipidated Pepstatin A. Pharmaceutics. 2023 Oct 14;15(10):2464.

30. Fischer M, Grundke I, Sohr S, Quaas M, Hoffmann S, Knörck A, et al. p53 and cell cycle dependent transcription of kinesin family member 23 (KIF23) is controlled via a CHR promoter element bound by DREAM and MMB complexes. PLoS One. 2013;8(5):e63187.

31. Yang X, Wang H, Zhang L, Yao S, Dai J, Wen G, et al. Novel roles of karyopherin subunit alpha 2 in hepatocellular carcinoma. Biomedicine & Pharmacotherapy. 2023 Jul 1;163:114792.

32. Lonati L, Barbieri S, Guardamagna I, Ottolenghi A, Baiocco G. Radiation-induced cell cycle perturbations: a computational tool validated with flow-cytometry data. Sci Rep. 2021 Jan 13;11(1):925.

33. Kaminskyy VO, Hååg P, Novak M, Végvári Á, Arapi V, Lewensohn R, et al. EPHA2 Interacts with DNA-PKcs in Cell Nucleus and Controls Ionizing Radiation Responses in Non-Small Cell Lung Cancer Cells. Cancers. 2021 Jan;13(5):1010.

34. Bailis JM, Forsburg SL. MCM proteins: DNA damage, mutagenesis and repair. Curr Opin Genet Dev. 2004 Feb;14(1):17–21.

35. Wu N, Yu H. The Smc complexes in DNA damage response. Cell Biosci. 2012 Feb 27;2:5.

36. Finney AC, Scott ML, Reeves KA, Wang D, Alfaidi M, Schwartz JC, et al. EphA2 signaling within integrin adhesions regulates fibrillar adhesion elongation and fibronectin deposition. Matrix Biol. 2021 Sep;103–104:1–21.

37. Pecci A, Ma X, Savoia A, Adelstein RS. MYH9: Structure, functions and role of non-muscle myosin IIA in human disease. Gene. 2018 Jul 20;664:152–67.

38. Li G, Jiang H, Wang L, Liang T, Ding C, Yang M, et al. The role of PALLD-STAT3 interaction in megakaryocyte differentiation and thrombocytopenia treatment. Haematologica. 2024 May 30;109(11):3693–704.

39. Simon NE, Schwacha A. The Mcm2-7 Replicative Helicase: A Promising Chemotherapeutic Target. Biomed Res Int. 2014;2014:549719.

40. Babel L, Grunewald M, Lehn R, Langhans M, Meckel T. Direct evidence for cell adhesion-mediated radioresistance (CAM-RR) on the level of individual integrin β1 clusters. Sci Rep. 2017 Jun 13;7(1):3393.

41. Huber RJ, Kim WD, Wilson-Smillie MLDM. Mechanisms regulating the intracellular trafficking and release of CLN5 and CTSD. Traffic. 2024 Jan;25(1):e12925.

42. Liu L, He L, Li W, Zhao T, Li G, Xiu X, et al. The miR-4306/IGF2R axis modulates the lung adenocarcinoma response to irradiation in vitro and in vivo. Transl Lung Cancer Res. 2021 Dec;10(12):4494–510.

43. Zheng W, Chen Q, Wang C, Yao D, Zhu L, Pan Y, et al. Inhibition of Cathepsin D (CTSD) enhances radiosensitivity of glioblastoma cells by attenuating autophagy. Mol Carcinog. 2020 Jun;59(6):651–60.

44. Zhang H, Song Q, Shang K, Li Y, Jiang L, Yang L. Tspan protein family: focusing on the occurrence, progression, and treatment of cancer. Cell Death Discov. 2024 Apr 22;10(1):1–14.

45. Zhang Y, Wang C, Xu Y, Su H. Tetraspanin 3 promotes NSCLC cell proliferation via regulation of β1 integrin intracellular recycling. Cellular & Molecular Biology Letters. 2024 Sep 27;29(1):124.

46. Liang LY, Patel O, Janes PW, Murphy JM, Lucet IS. Eph receptor signalling: from catalytic to non-catalytic functions. Oncogene. 2019 Sep;38(39):6567–84.

47. Kaplan N, Wang S, Wang J, Yang W, Ventrella R, Majekodunmi A, et al. Ciliogenesis and autophagy are coordinately regulated by EphA2 in the cornea to maintain proper epithelial architecture. Ocul Surf. 2021 Jul;21:193–205.

48. Tandon M, Vemula SV, Mittal SK. Emerging strategies for EphA2 receptor targeting for cancer therapeutics. Expert Opin Ther Targets. 2011 Jan;15(1):31–51.

49. Waller V, Tschanz F, Winkler R, Pruschy M. The role of EphA2 in ADAM17- and ionizing radiation-enhanced lung cancer cell migration. Front Oncol. 2023;13:1117326.

50. Graves PR, Din SU, Ashamalla M, Ashamalla H, Gilbert TSK, Graves LM. Ionizing radiation induces EphA2 S897 phosphorylation in a MEK/ERK/RSK-dependent manner. Int J Radiat Biol. 2017 Sep;93(9):929–36.

51. Gong S, Li Y, Lv L, Men W. Restored microRNA-519a enhances the radiosensitivity of non-small cell lung cancer via suppressing EphA2. Gene Ther. 2022 Nov;29(10):588–600.

52. Amato KR, Wang S, Hastings AK, Youngblood VM, Santapuram PR, Chen H, et al. Genetic and pharmacologic inhibition of EPHA2 promotes apoptosis in NSCLC. J Clin Invest. 2014 May 1;124(5):2037–49.

53. Perez-Riverol Y, Bai J, Bandla C, García-Seisdedos D, Hewapathirana S, Kamatchinathan S, et al. The PRIDE database resources in 2022: a hub for mass spectrometry-based proteomics evidences. Nucleic Acids Research. 2022 Jan 7;50(D1):D543–52.

