## Supplementary for "Proteome changes associated with effect of high-dose single-fractionation radiation on lung adenocarcinoma cell lines"

**Supplementary Data**

*Supplementary Table Legends*

Table S1. List of differentially expressed proteins in irradiated vs non-irradiated cells

In all tabs, FDR stands for the false discovery rate and log2 (fold change) for the binary logarithm of the average label-free intensity ratio measured for each listed protein in the irradiated *vs* non-irradiated sample. Positive log2 (fold change) values correspond to proteins upregulated following irradiation and negative log2 (fold change) values correspond to proteins downregulated following irradiation.

- Tab HCC_2D_10Gy_vs_HCC_2D_NT: comparison of irradiated *vs* non-irradiated HCC-44 cells grown adherently;
- Tab A549_2D_10Gy_vs_A549_2D_NT: comparison of irradiated *vs* non-irradiated A549 cells grown adherently;
- Tab A549_sph_10Gy_vs_A549_sph_NT: comparison of irradiated *vs* non-irradiated A549 cells grown as spheroids.

Table S2. Cellular pathways significantly up- or downregulated in irradiated vs non-irradiated cells

The analysis was carried out using the online platform g:Profiler (https://biit.cs.ut.ee/gprofiler/gost) based on the differentially expressed proteins listed in Table S1.

- Tab HCC_2D_up after irr: pathways upregulated in irradiated *vs* non-irradiated HCC-44 cells grown adherently;
- Tab HCC_2D_down after irr: pathways downregulated in irradiated *vs* non-irradiated HCC-44 cells grown adherently;
- Tab A549_2D_up after irr: pathways upregulated in irradiated *vs* non-irradiated A549 cells grown adherently;
- Tab A549_2D_down after irr: pathways downregulated in irradiated *vs* non-irradiated A549 cells grown adherently;
- Tab A549_sph_down after irr: pathways down regulated in irradiated *vs* non-irradiated A549 cells grown as spheroids.

No pathways were found to be significantly upregulated in irradiated *vs* non-irradiated A549 cells grown as spheroids.

*Supplementary Figures*

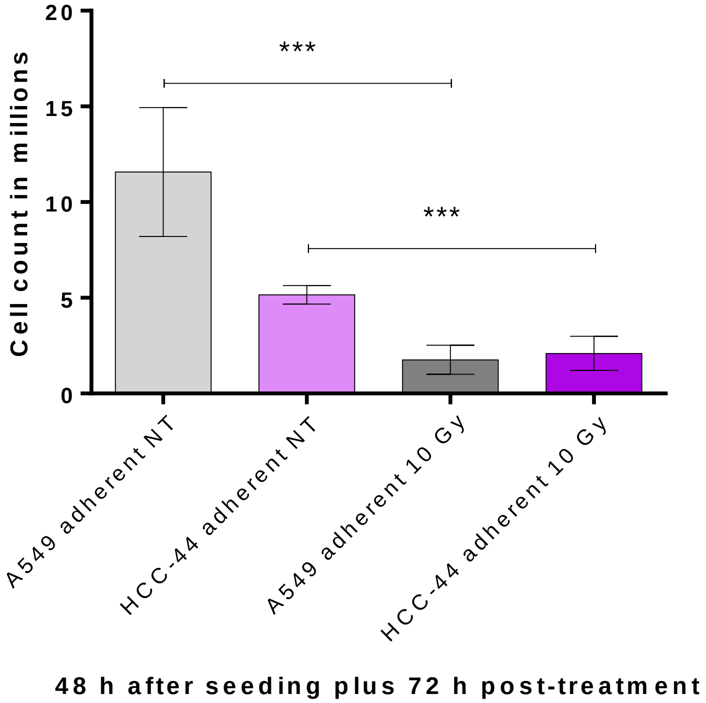


Figure S1. Total cell count in the irradiated and non-irradiated samples before preparation of pellets for the mass-spectrometry. The graph shows average cell count ± standard deviation for three independent experiments (cells grown adherently prior to trypsinization). Statistical significance of difference between the irradiated (10 Gy) and non-irradiated (NT) samples: *** corresponds to $P < 0.001$.

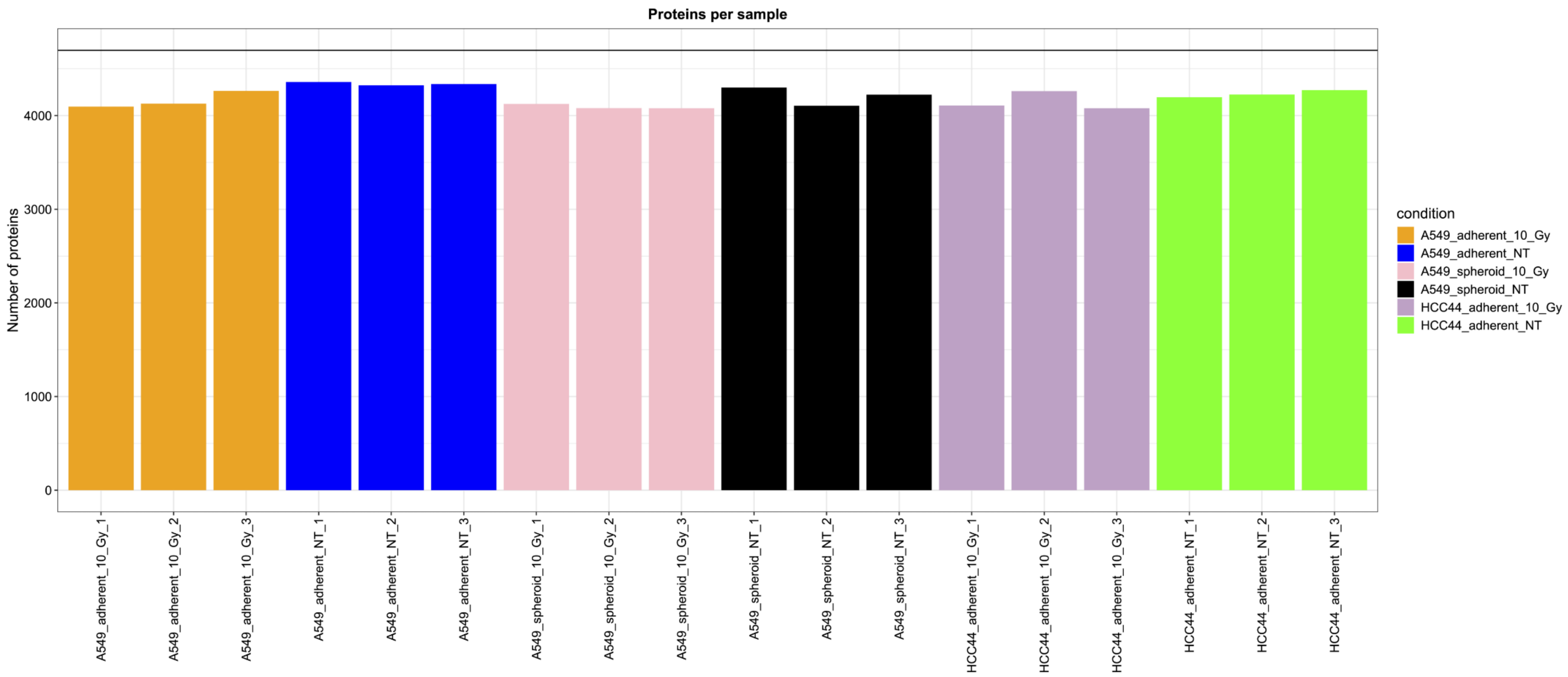


Figure S2. Bar plots depicting the number of proteins per sample as identified by the mass-spectrometry. Each vertical bar represents an individual sample. The treatment conditions are listed on the right and below each column; NT stands for not irradiated cells and 10 Gy for irradiated cells.

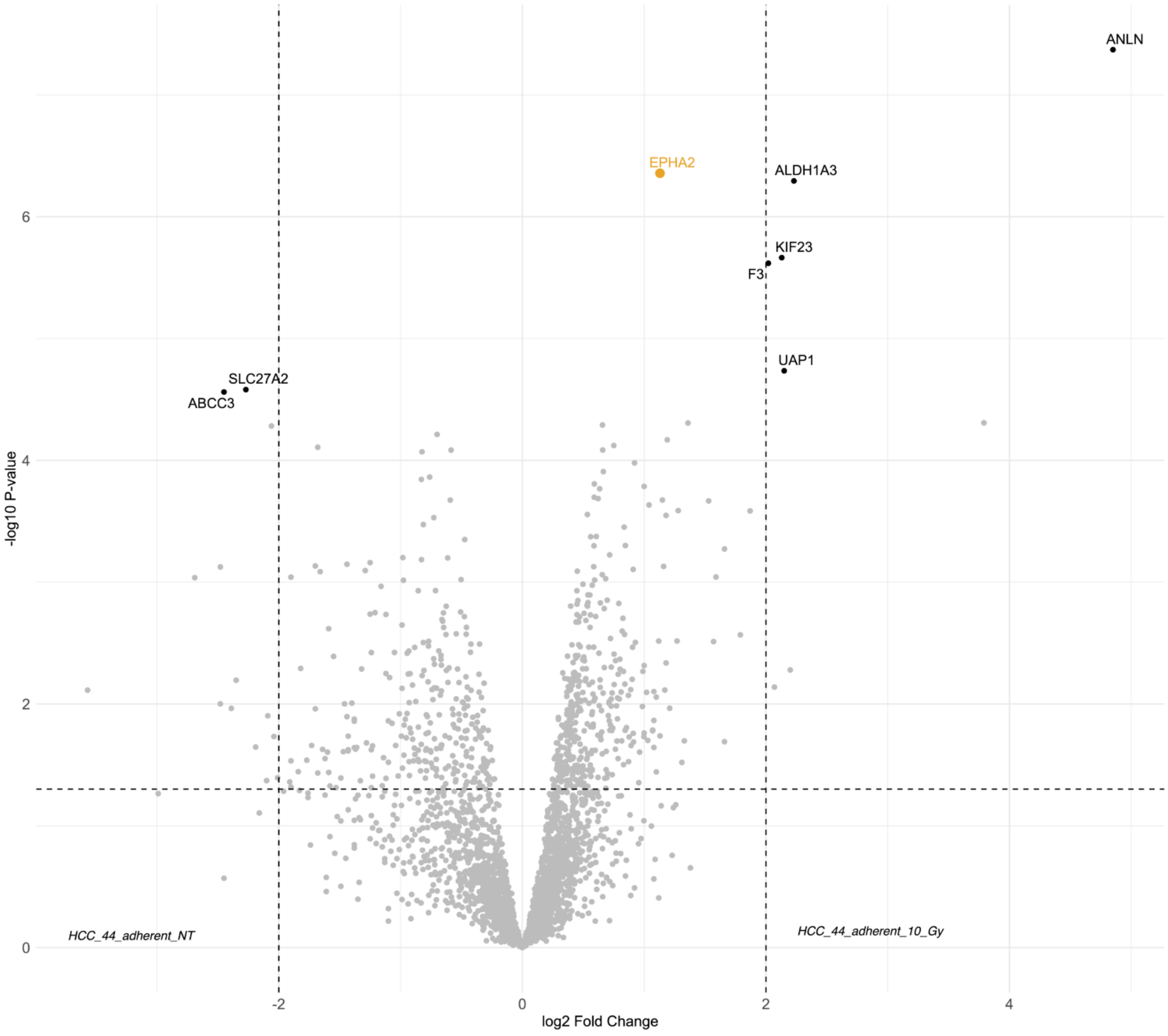


Figure S3. Volcano plot showing differentially expressed proteins in the irradiated *vs* non-irradiated adherent HCC-44 cells. Top hits are marked with the name labels, and a target of interest chosen for the validation in the subsequent steps (ephrin type-A receptor 2, EPHA2) is shown in orange.

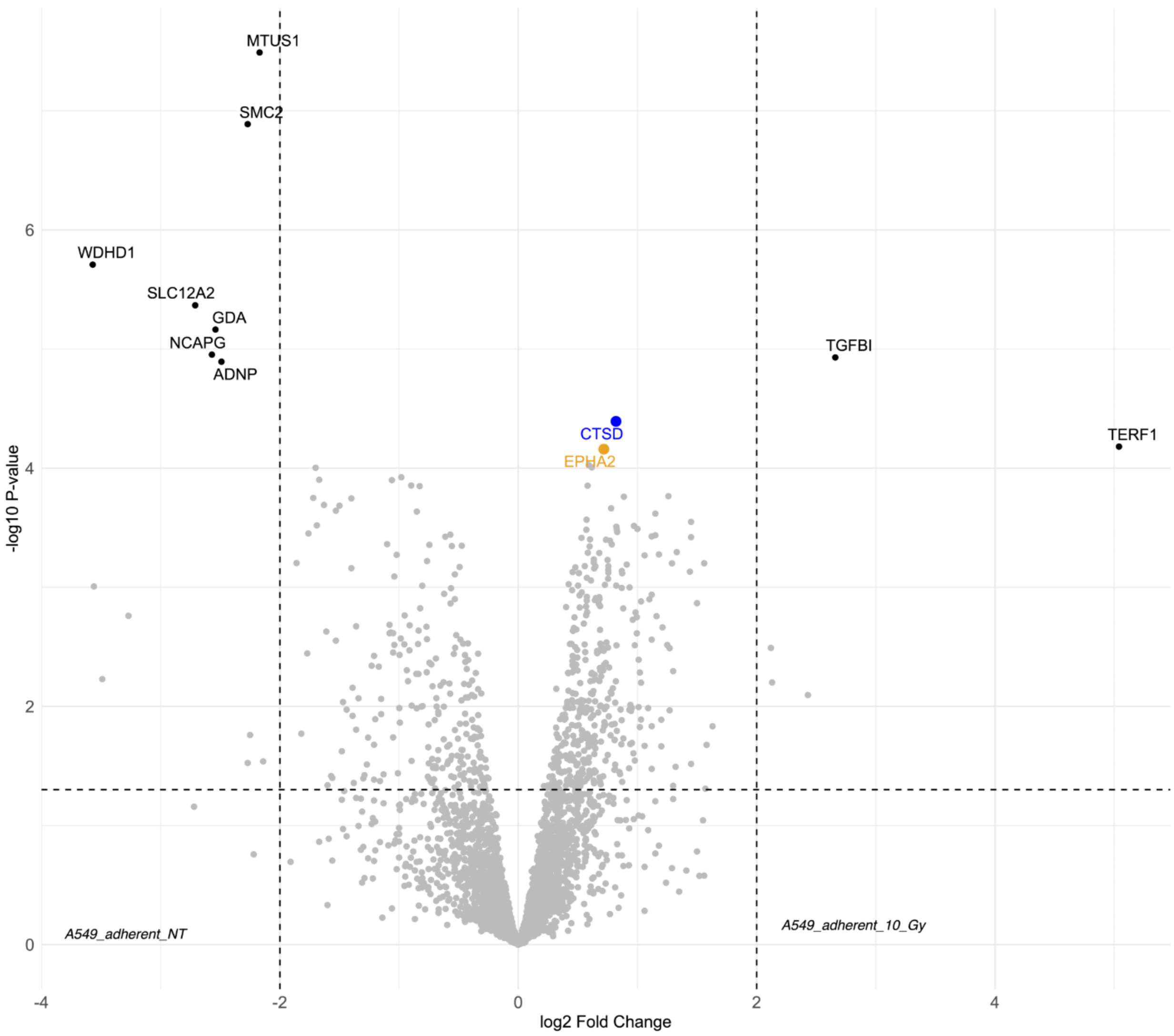


Figure S4. Volcano plot showing differentially expressed proteins in the irradiated *vs* non-irradiated adherent A549 cells. Top hits are marked with the name labels, and the targets of interest chosen for the validation in the subsequent steps are shown in orange (ephrin type-A receptor 2, EPHA2) or blue (cathepsin D, CTSD).

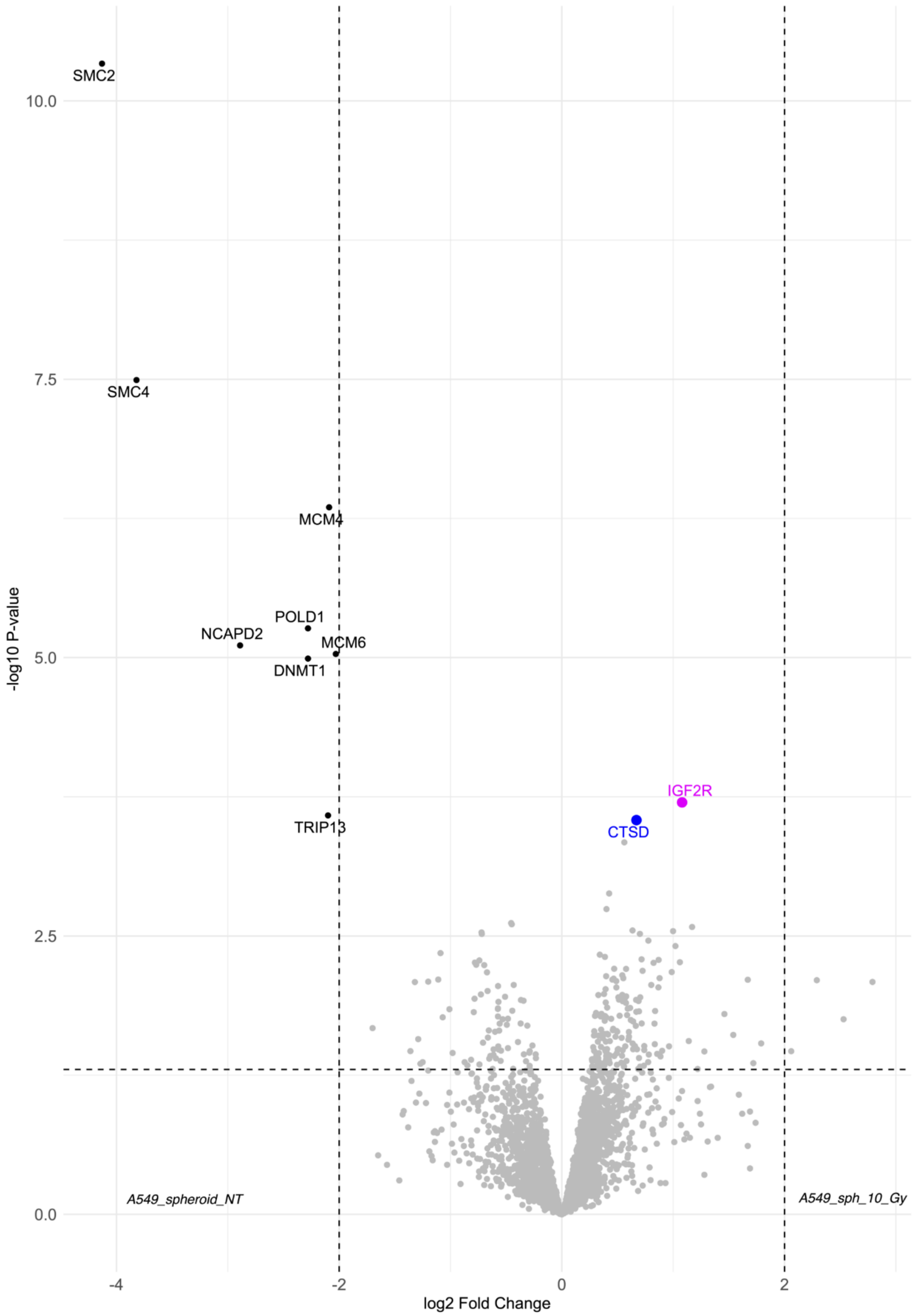


Figure S5. Volcano plot showing differentially expressed proteins in the irradiated *vs* non-irradiated A549 spheroids. Top hits are marked with the name labels, and the targets of interest chosen for the validation in the subsequent steps are shown in blue (cathepsin D, CTSD) or pink (insulin-like growth factor 2 receptor, IGF2R).

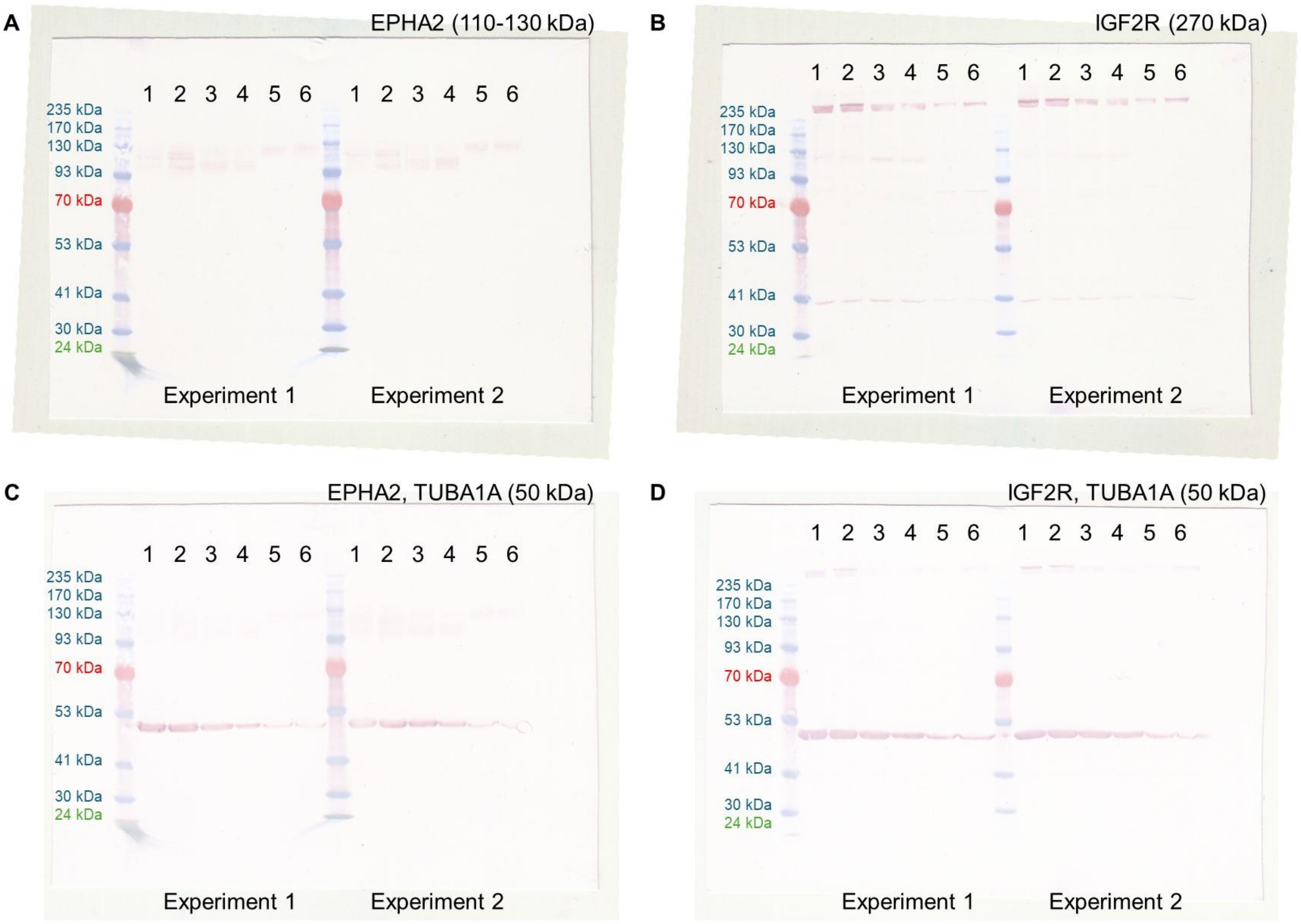


Figure S6. Western blot membranes from experiments 1-2. A and C, EPHA2 staining; B and D, IGF2R staining. In C and D, TUBA1A staining was performed following the staining of EPHA2 or IGF2R and reactivation of the membrane by methanol. Lane 1 corresponds to non-irradiated HCC-44 cells grown adherently, lane 2 to irradiated HCC-44 cells grown adherently, lane 3 to non-irradiated A549 cells grown adherently, lane 4 to irradiated A549 cells grown adherently, lane 5 to non-irradiated A549 cells grown as spheroids, lane 6 to irradiated A549 cells grown as spheroids. For better visualization, the image contrast in panels A and B was reduced.

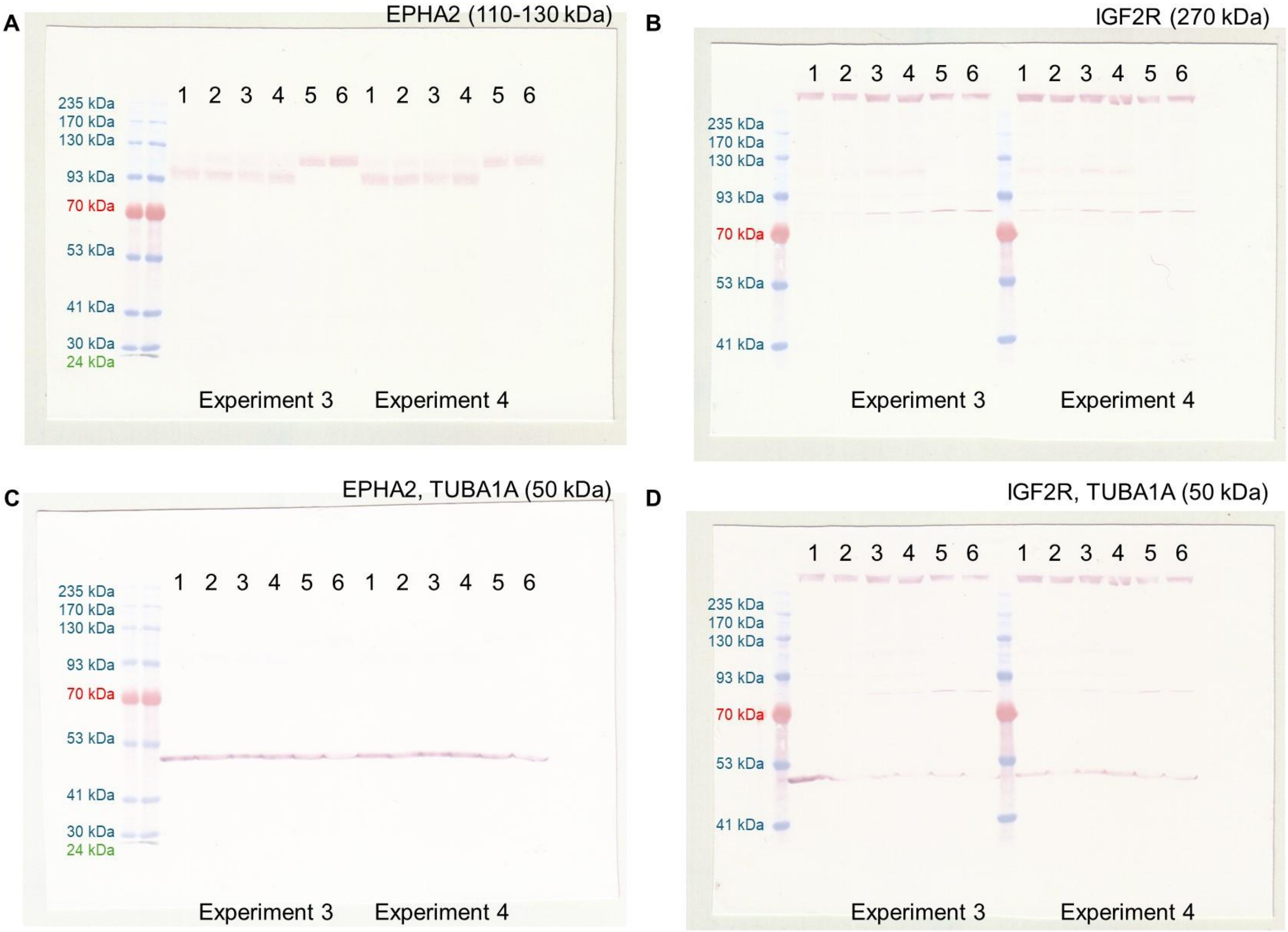


Figure S 7. Western blot membranes from experiments 3-4. A and C, EPHA2 staining; B and D, IGF2R staining. In C and D, TUBA1A staining was performed following the staining of EPHA2 or IGF2R and reactivation of the membrane by methanol. Lane 1 corresponds to non-irradiated HCC-44 cells grown adherently, lane 2 to irradiated HCC-44 cells grown adherently, lane 3 to non-irradiated A549 cells grown adherently, lane 4 to irradiated A549 cells grown adherently, lane 5 to non-irradiated A549 cells grown as spheroids, lane 6 to irradiated A549 cells grown as spheroids. For better visualization, the image contrast in panels A, B and D was reduced.

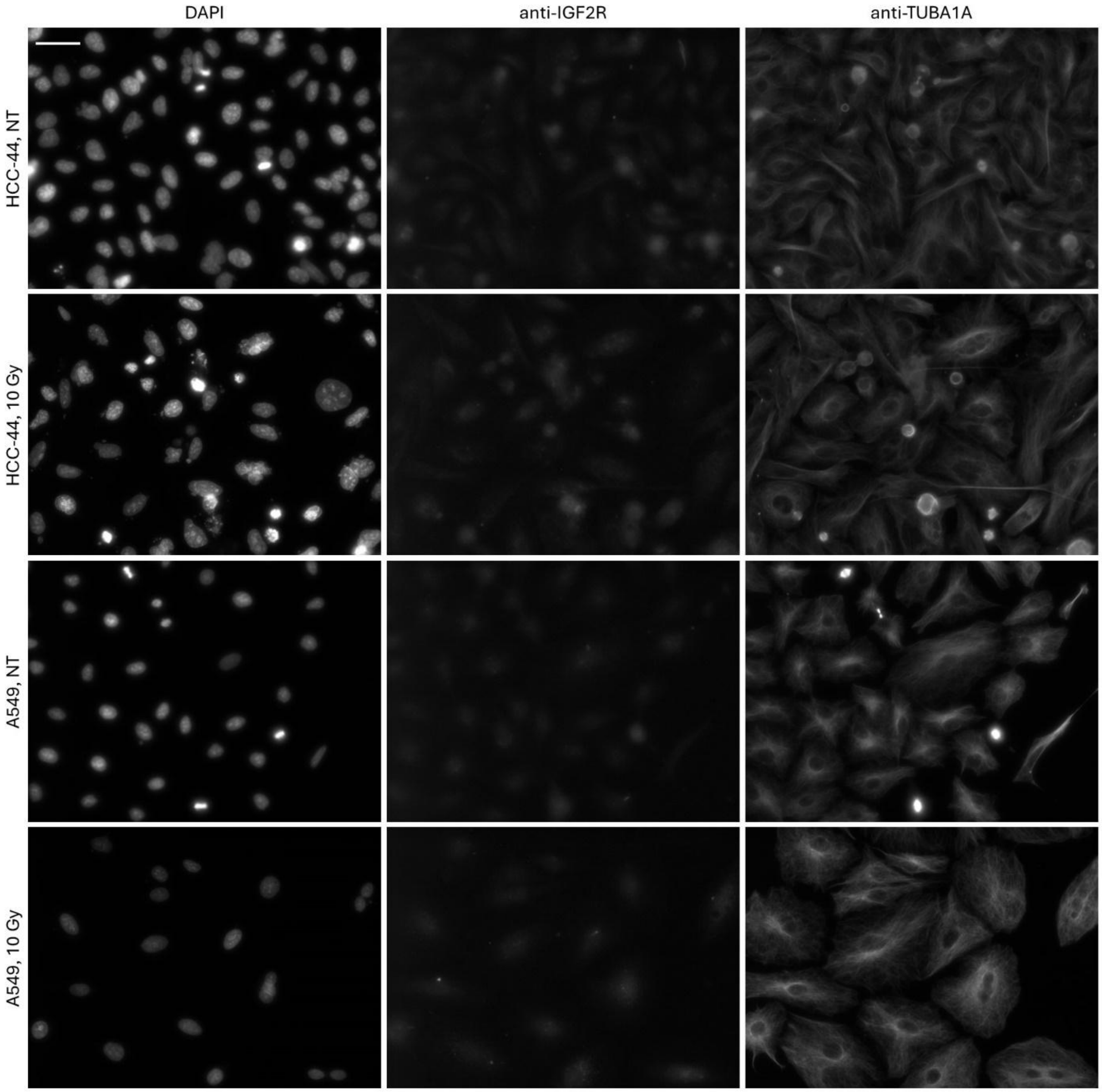


Figure S8. Representative examples of IGF2R immunostaining in irradiated *vs* non-irradiated adherent fixed cells. Data from a single experiment is shown; the channels (nuclear stain DAPI, protein of interest, and TUBA1A) are listed above the images; the cell lines and treatment conditions are listed on the left (NT stands for not irradiated). Scale bar (top left): 50 μm.

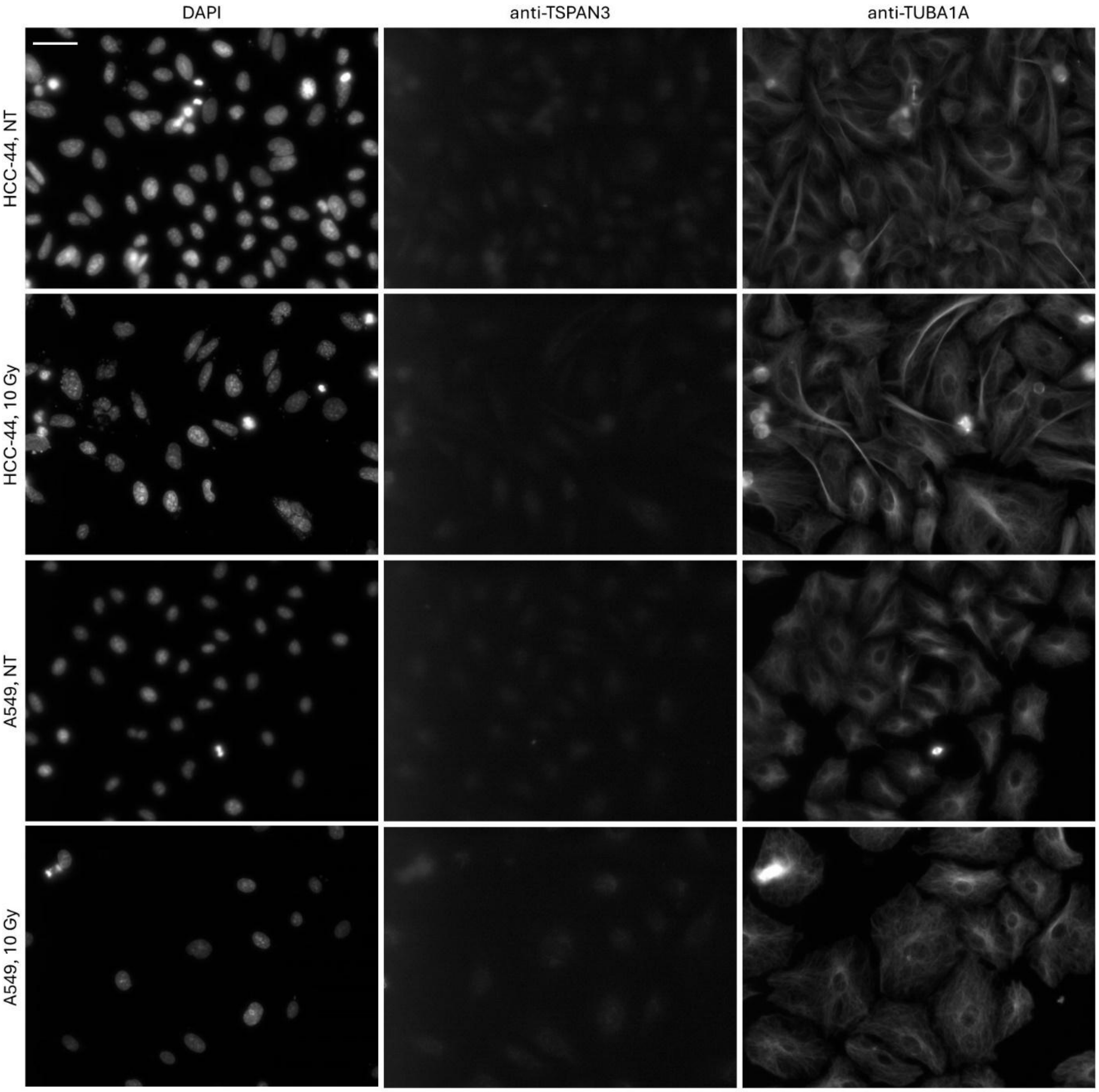


Figure S9. Representative examples of TSPAN3 immunostaining in irradiated *vs* non-irradiated adherent fixed cells. Data from a single experiment is shown; the channels (nuclear stain DAPI, protein of interest, and TUBA1A) are listed above the images; the cell lines and treatment conditions are listed on the left (NT stands for not irradiated). Scale bar (top left): 50 μm. For better visualization, the brightness in anti-TSPAN3 channel was increased by 20% and the contrast was reduced by 40%; the quantification data shown in the main text corresponds to the non-modified images.

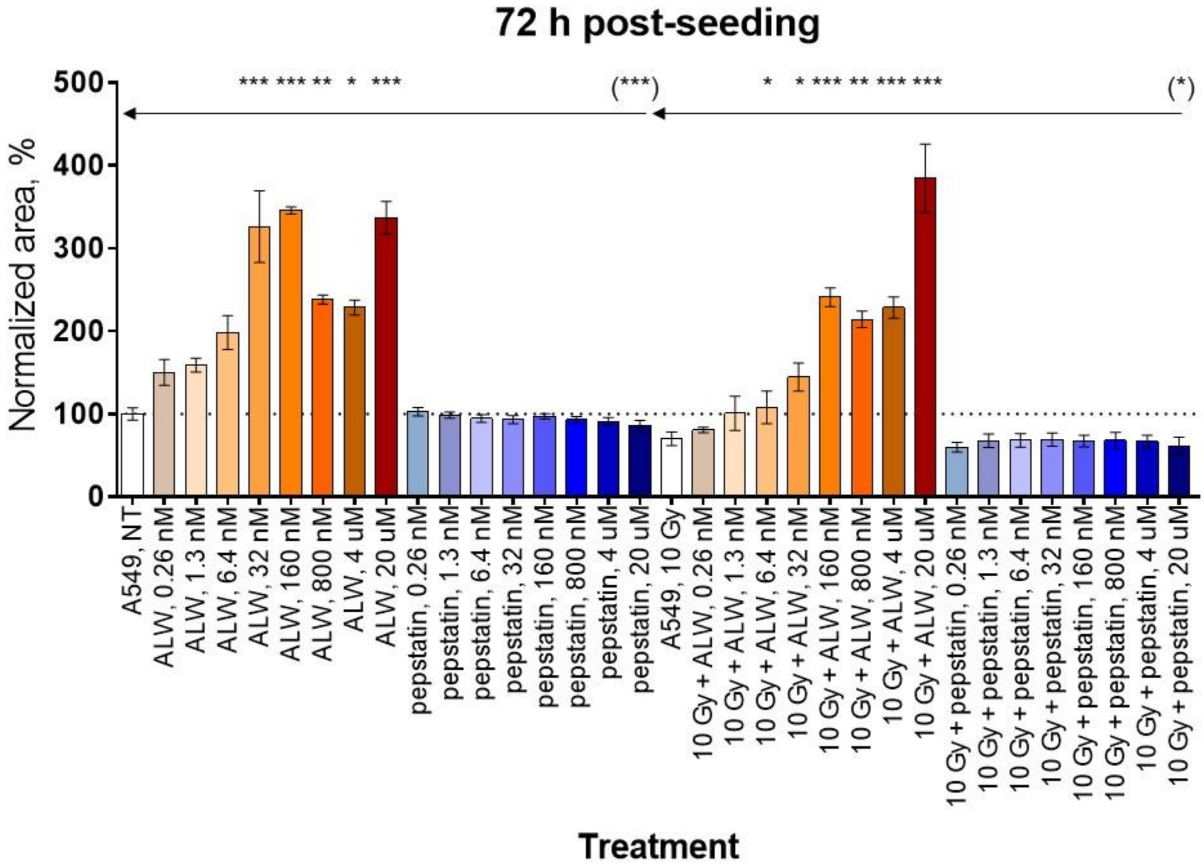


Figure S10. Pooled normalized area of A549 spheroids at 72 h post-seeding (N = 3); the treatment conditions are listed below the graphs. Each column represents average ± standard deviation. Statistical significance of the grouped comparisons (Kruskal-Wallis test): *** corresponds to $P < 0.001$; ** corresponds to $P < 0.01$; * corresponds to $P < 0.05$; only significant comparisons are shown; arrows indicate the treatments (non-treated A549 or irradiated A549) to which other treatments are compared. Statistical significance of paired comparisons (Mann-Whitney test, only carried out for pepstatin A treatment *versus* cells not treated with inhibitors): (***) corresponds to $P < 0.001$; (*) corresponds to $P < 0.05$; only significant comparisons are shown.
